# Mechanical Cues Regulate Cargo Sorting and Export at the Golgi

**DOI:** 10.1101/2025.09.09.675067

**Authors:** Greta Serafino, Stefania Forciniti, Edoardo Scarpa, Antonio Ranieri, Lucia Santorelli, Gabriella De Blasi, Sara Costantini, Eunjoo Jun Lee, Antonio Galeone, Alessia Calcagnì, Marinella Pirozzi, Loretta L. del Mercato, Rossella Venditti, Giovanni D’Angelo, Seetharaman Parashuraman, Tiziano Verri, Giuseppe Gigli, Elizabeth S. Sztul, Paolo Grumati, Loris Rizzello, Domenico Russo, Riccardo Rizzo

## Abstract

The secretory pathway is a sophisticated endomembrane machinery designed to transport and deliver proteins and lipids to intracellular organelles and the extracellular space. While the molecular components of the secretory pathway are well understood, less is known about their regulation, especially by mechanical cues. Here, we report that substrate stiffness stimulates conventional secretion. We have unravelled a molecular pathway that links a mechanical cue through Src and FAK kinases to promote the trafficking of secretory proteins out of the Golgi apparatus and prevent their post-Golgi lysosomal degradation. Phosphoproteomic analysis revealed the Golgi-specific Brefeldin A resistance factor 1 (GBF1) as a key downstream mechano-responsive regulator, whose phosphorylation state orchestrates post-Golgi cargo sorting, directing proteins either toward secretion or to lysosomes. Finally, we identified AMPK as a stiffness-dependent upstream regulator of GBF1 phosphorylation. Together, our data reveal a molecular regulatory loop in which matrix stiffness positively regulates cellular secretion via the Src-FAK-AMPK-GBF1 axis, which can have relevant medical implications in conditions like cancer and fibrosis and their treatment.

## 1. Introduction

Mammalian cells experience social life by interacting with other cells and the extracellular environment via signalling and structural proteins, hormones, receptors, etc^[1,2]^. Most of these proteins are synthesised by the secretory pathway, an endomembrane system designed to transport, process and deliver proteins and lipids to the endolysosomal system or the plasma membrane (PM). Cargoes following the secretory pathway are synthesised in the endoplasmic reticulum (ER) and transported to the Golgi complex via COPII carriers ^[3]^. At the Golgi, proteins are processed ^[4–8]^ and then sorted at the trans-Golgi network (TGN) to their final destination^[9]^. As such, the secretory pathway represents an intra- and intercellular communication platform that plays a key role in diseases including cancer and fibrosis ^[9,10]^. While numerous components of the molecular machinery of the secretory pathway are well understood ^[11]^, less is known about their regulation, with most of the existing knowledge focused on biochemical stimuli (i.e. nutrients, hormones, hypoxia, inflammatory cytokines, etc.).

Such regulation is known to affect both the functional organisation of the secretory pathway and the secretory output, in response to the secretion demands imposed by these perturbations ^[12]^. Initial evidence supporting this hypothesis arises from experiments demonstrating the impact of growth factors and second messengers on Golgi to PM transport ^[13]^. Other stimuli, such as hypoxia and glucose levels, have been demonstrated to affect the transport of proteins from the ER to the Golgi apparatus ^[14]^, as well as the fusion of secretory granules with the PM ^[15]^, respectively. Moreover, intracellular auto-regulatory circuits have been identified and shown to regulate various trafficking steps and protein synthesis in response to cargo load, maintaining cellular homeostasis and preventing harmful fluctuations in secretion ^[16–18]^. These examples demonstrate how biochemical stimuli can regulate different steps of transport.

Besides chemical stimuli, cells within tissues are also exposed to a variety of mechanical stimuli, which prompt cells to make decisions affecting their phenotype ^[19–22]^. These include compression, stiffness, traction forces or shear stress, which are sensed and transduced to elicit a biological cellular response ^[23]^. The PM, nucleus and cytoskeleton serve as receptors and mediators of mechanical signalling and have been the focus of much research in this area ^[24]^.

Nevertheless, the extent to which mechanical cues affect the secretory pathway is currently poorly understood. Recent evidence in this new field demonstrated that mechanical tension, induced by hypotonic swelling or biaxial stretch, stimulates ER-Exit Sites (ERES) number and enhanced the ER to Golgi transport of Mannosidase II via Rac1 ^[25]^. Similarly, the matrix stiffness, commonly altered in fibrotic diseases and known to regulate several hallmarks of cancer^[26–28]^, has been demonstrated to increases PM tension, a process promoting Tissue Inhibitor of Metalloproteinases 1 (TIMP1) exocytosis via Caveolin1 and Dynamin2 ^[29]^.

Following up on this thread, we discovered a novel regulatory paradigm that controls protein secretion influenced by mechanical cues. We found that extracellular matrix stiffness (ECM), a mechanical property reflecting the ECM resistance to deformation, positively regulates protein transport from the Golgi to the PM for secretion. This occurs by inhibiting lysosomal trafficking and degradation in a FAK-dependent manner; and controls the transport of cargo towards PM via Src. By exploring changes in protein phosphorylation during cellular mechanosensing using phosphoproteomics, we found Golgi-specific Brefeldin A resistance factor 1 (GBF1) as a downstream key mechanotransducer regulating post-Golgi sorting. We further established that AMP-activated protein kinase (AMPK) functions as an upstream regulator of GBF1 phosphorylation in a stiffness-dependent manner. Specifically, phosphorylation of GBF1 at serine 1318 (S1318) promotes cargo protein degradation via the lysosomal pathway, thereby attenuating secretion. This proves GBF1 as a one key regulator of protein secretion triggered by mechanical cues, and represents a novel target for antifibrotic treatment in conditions such as cancer and fibrosis.

## 2. Results

### 2.1. Substrate stiffness correlates with protein secretion

We started exploring the link between substrate stiffness and conventional secretion through a broad investigation of cell surface components aimed at quantifying the deposition of nascent proteins and glycoproteins on PM under substrate stiffness perturbation. To this end, we cultured HeLa cells on stiff fibronectin-coated plastic (here referred as >Giga Pascal, GPa) or soft silicone plates (CytoSoft 0.5 kilopascal (kPa) and 8 kPa), respectively (Figure 1A). The selected stiffness values align with established physiological and pathological stiffness ranges, representing soft healthy tissues (0.2-2 kPa, reflecting the normal mechanical properties of healthy tissues), pathological fibrotic and cancerous microenvironments (8-12 kPa), and rigid mineralized bone tissue (typically in the range of gigapascal, GPa), providing a comprehensive model for diverse cellular responses ^[30–32]^. We performed a metabolic labelling assay, to quantify protein secretion, by administrating a modified azide-methionine analogue (azidohomoalanine, AHA), or the sugar analogue azide-mannose, (ManNAz) followed by a bio-orthogonal strain-promoted cyclo-addition with fluorophore-modified cyclooctyne (DBCO-Cy5 ^[33,34]^). The assay was performed in non-permeabilized cells, allowing selective detection of surface-exposed proteins only. We found that increased rigidity of the substrate correlates with increased deposition of methionine-(Figure 1B, D) or mannose-(Figure 1C, E) containing transmembrane proteins present at the PM. To exclude that differences in surface protein deposition were due to substrate-dependent variations in labelling efficiency, we repeated the assay with a short 1 hour pulse with AHA or ManNAz. Cells were then fixed and permeabilized to detect intracellular AHA- or ManNAz-labeled proteins with DBCO-Cy5. Total intracellular labelling was comparable across all substrates, indicating similar labelling efficiency and excluding it as a factor contributing to altered surface deposition (Figure S1A-D).

**Figure 1.**
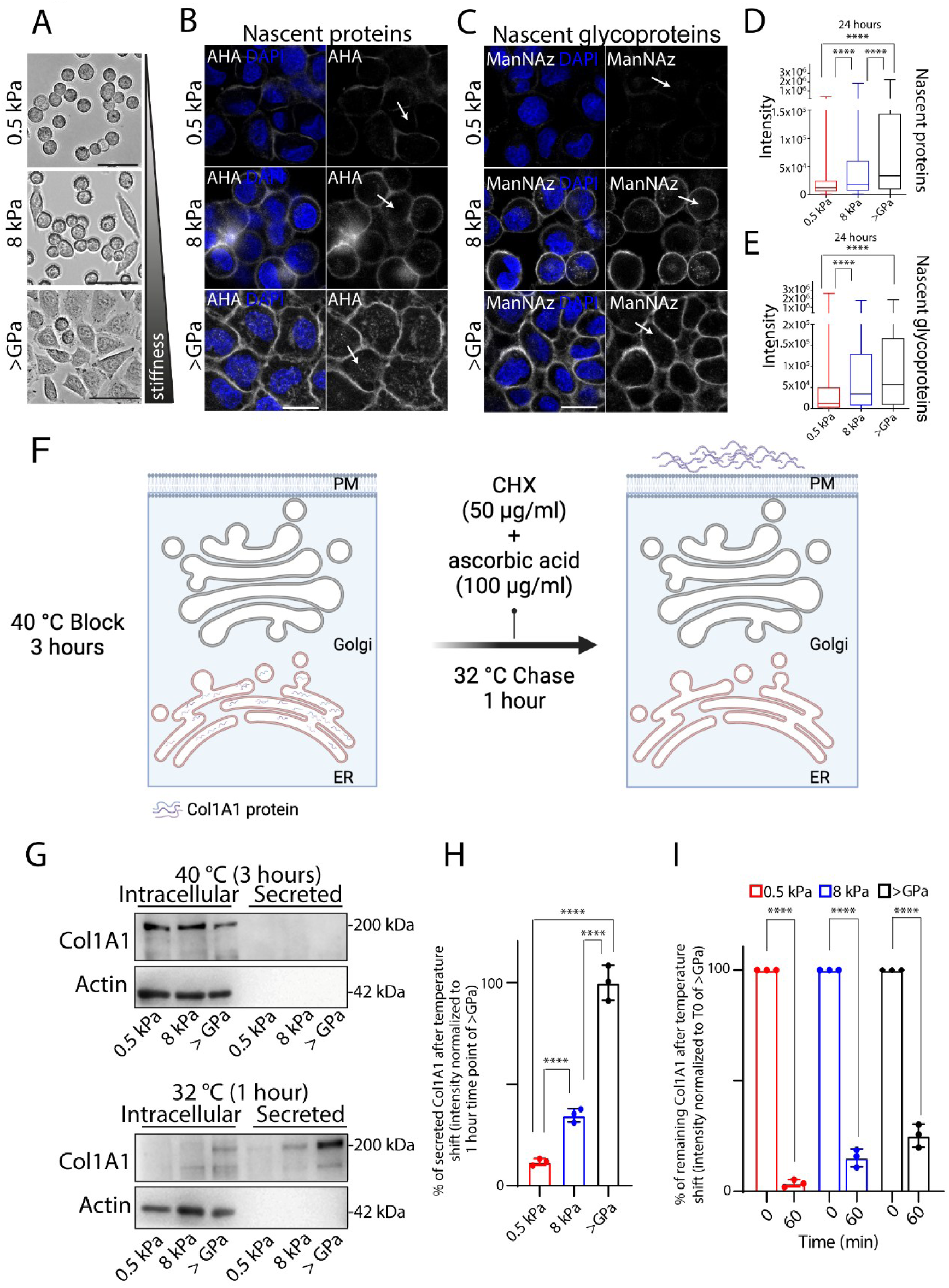
Substrate stiffness correlates with protein secretion. **A.** Bright-field images of HeLa cells cultured for 24 hours on indicated substrate rigidities (0.5 kPa; 8 kPa, >GPa). Scale bar, 100 μm. **B**, **C**. HeLa cells were grown on the indicated substrate rigidities coated with 10 μg/ml human fibronectin. The azide-modified methionine analogue (AHA, [B]) or a sugar analogue (ManNAz, [C]) was added to the culture media for 24 hours prior to proceed with bio-orthogonal strain-promoted cyclo-addition with fluorophore-modified cyclooctyne (DBCO-Cy5). Representative images of PM nascent proteins (AHA, [B]) and glycoproteins (ManNAz, [C]) are shown. DAPI (blue), AHA or ManNAz (white). Scale bar, 20 μm. **D**, **E**. Quantification of the PM fluorescence intensity of nascent proteins (D) and glycoproteins (E) of the experiments in (B) and (C) (data are means ± SEM derived from 2 biological replicates, in which cumulatively >540 cells have been quantified for each condition. ****p<0.0001 [One-way ANOVA]). **F**. Synchronisation protocol of procollagen I secretion (created with BioRender.com). BJ-5ta cells were incubated for 3 hours at 40°C, then shifted to 32°C for 1 hours in the presence of cycloheximide (CHX, 50 μg/ml) and ascorbic acid (100 μg/ml) (see Methods). **G**. Western blot analysis of collagen secretion (Col1A1 antibody was used to detect PC-I protein) in BJ-5ta fibroblasts grown on indicated substrate stiffness. Cells were grown on indicated substrate rigidities and after 24 hours the transport of Col1A1 was synchronized according to the ascorbic acid-CHX/40-32°C synchronization protocol. At each indicated time points cells were lysed and the cell lysate (intracellular) and the medium (secreted) were collected at indicated timepoint and analysed by SDS-PAGE (20% of the total intracellular lysate and 100% of the total secreted was loaded, respectively) followed by Western blot. **H, I**. Densitometric quantification of the secreted (H) and intracellular (I) Col1A1 of the blot in (G) is plotted (data are means ± SD of at least 3 independent experiments). ****p<0.0001, [Two-way ANOVA]).

Next, we aimed to evaluate the effect of substrate stiffness on soluble cargo proteins secretion. To this aim we cultured HeLa cells on substrates of varying rigidity (0.5 kPa, 8 kPa and >GPa) at 10°C for 3 hours to synchronize protein synthesis, and to load and accumulate newly synthesized proteins in the ER ^[35]^. This was followed by a temperature shift to 37°C in the presence of cycloheximide (CHX) to inhibit protein synthesis (see Methods, Figure S1E). After a 2 hours chase, we collected media from cells cultured on each substrate and performed a secretome profiling using an antibody-based protein array ^[36]^, which allowed the detection of a set of cargo proteins abundantly expressed in our HeLa cell system ^[2]^. Out of a total of 80 identified targets, the secretion levels of 14 proteins (CXCL5, G-CSF, GRO, CXCL1, IL-12, Angiogenin, IGFBP-2, IGFBP-3, NT-3, PLGF, TGF-β 2, TGF-β3, TIMP1 and TIMP2) increased with substrate stiffness (Figure S1F, G), while the majority remained unchanged between 0.5 kPa and 8 kPa, as revealed by densitometric quantification (Figure S1H).

Importantly, no differences in the endogenous mRNA levels of these proteins were detected across different rigidities, except for CXCL5 and CXCL1 (Figure S1I). This indicates that substrate stiffness does not affect the transcription of G-CSF, GRO, IL-12, Angiogenin, IGFBP2, IGFBP3, NT3, PLGF, TGFβ 2, TGFβ3, TIMP1 and TIMP2. In line with our findings, TIMP1 exocytosis was shown to be regulated by substrate stiffness in a transcription-independent manner ^[29,37]^.

We further extended our analysis of an additional soluble cargo by tracking the transport of Procollagen-I (PC-I), a key component of the extracellular matrix (ECM) whose dysregulation is associated with fibrotic diseases^[38]^. BJ-5ta cells, a human immortalized fibroblast cell line, were cultured on substrates of varying stiffness (0.5 kPa, 8 kPa, and >GPa) and incubated at 40 °C for 3 hours to accumulate PC-I in the ER due to impaired protein folding. The temperature was then shifted to 32 °C, allowing PC-I to achieve its proper conformation and progress through the secretory pathway^[39]^ (see Methods and Figure1F). Western blot analysis revealed that 1 hour after PC-I release from the ER, intracellular cargo levels significantly decreased across all substrate rigidities. Surprisingly, in cells cultured on 0.5 kPa and 8 kPa substrates, this reduction did not correspond to a proportional increase in collagen secretion into the culture media (Figure 1G-I). These findings suggest that substrate stiffness might modulates both PC-I degradation and secretion.

Collectively, our results suggest that stiffness could affect the turnover (i.e., synthesis and-or degradation) and trafficking of nascent proteins, glycoproteins and PC-I. These observations prompted us to investigate the underlying mechanisms further.

### 2.2. Substrate stiffness regulates secretion by controlling the trafficking of secretory proteins

To evaluate the impact of substrate rigidity on conventional protein trafficking, we employed a well-established assay for measuring constitutive secretion based on HeLa cells stably expressing a human growth hormone construct (hGH-FM2-GFP) as a cargo reporter (HeLa-GH), whose transport can be synchronized in a ligand-dependent manner ^[40]^ (see synchronisation protocol, Figure 2A). To note, hGH-FM2-GFP bears a furin-cleavage site, that enables the monitoring of the trafficking events as the construct reach the TGN (see the 53 kDa band in Figure 2B). It should be emphasized that the presence of unprocessed construct (see the 75 kDa band in Figure 2B) in the media may be due to saturation of the furin cleavage ^[40]^ and that furin levels and localization can vary both between cells and across experiments ^[40,41]^. For these reasons, we do not specifically focus on furin-dependent hGH-FM2-GFP processing and instead consider all secreted forms, both processed and unprocessed (75 kDa and 53 kDa bands), in our analysis.

**Figure 2.**
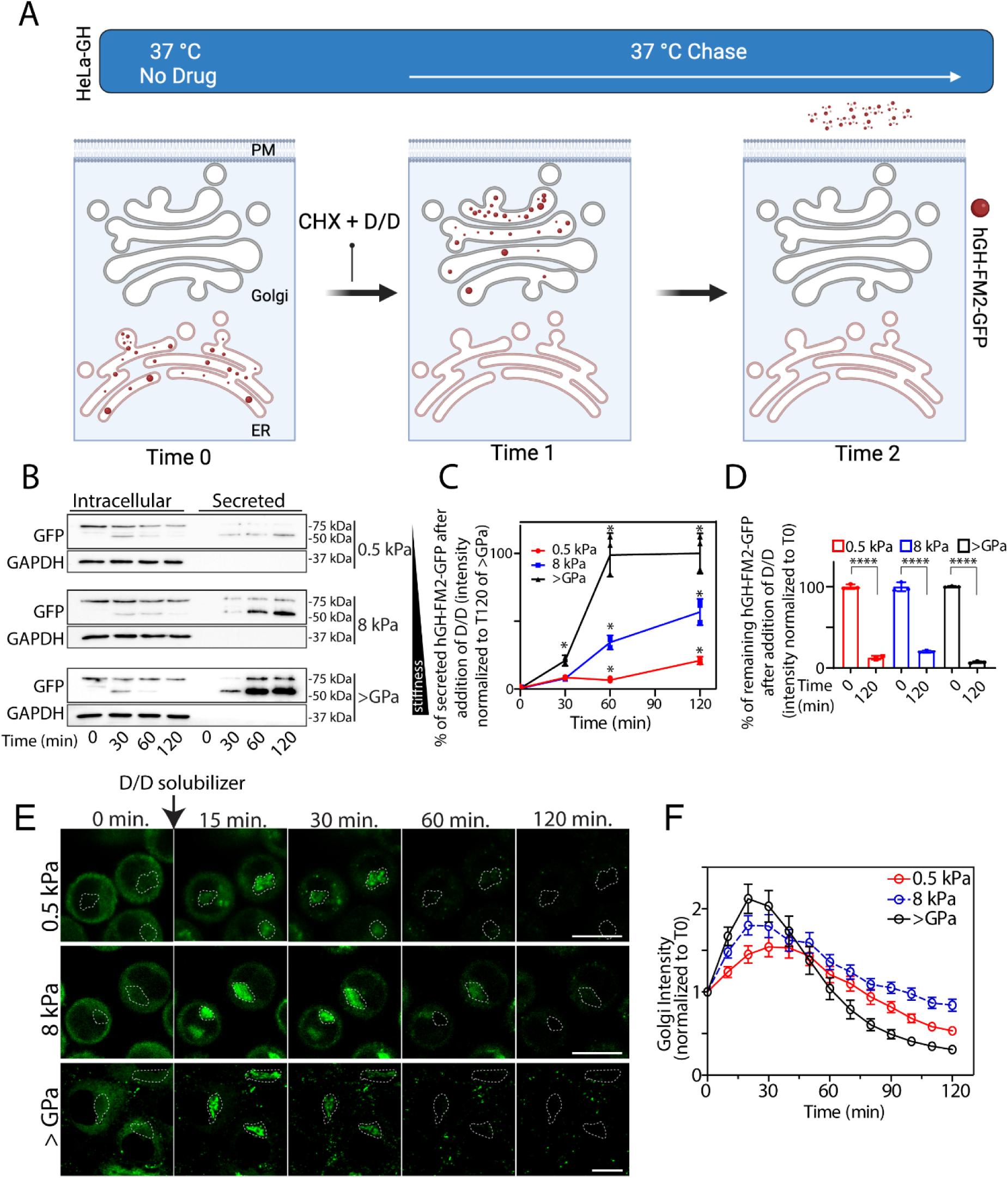
Substrate stiffness regulates secretion by controlling the trafficking of secretory proteins. **A**. Experimental scheme to probe secretion of the human growth hormone (hGH-FM2-GFP) cargo (created with BioRender.com). HeLa cells stably expressing hGH-FM2-GFP (HeLa-GH) were grown on 2D silicone with defined elastic modulus (0.5 kPa, 8 kPa and >GPa). After 24 hours, cells were subjected to ligand (D/D solubilizer, D/D, 1 μ_M_) to induce hGH-FM2-GFP release from the ER, in the presence of cycloheximide (CHX, 50 μg/ml), and sampled at the indicated time points. **B**. Cells were treated as in (A) and the cell lysate (intracellular) and the medium (secreted) were collected at indicated timepoint and analysed by SDS-PAGE (20% of the total intracellular lysate and 100% of the total secreted was loaded, respectively) followed by Western blot. **C**. Densitometric quantification of the blot in (B) is plotted (data are means ± SEM of at least 2 independent experiments. *p<0.05, [Two-way ANOVA]). **D**. Densitometric quantification of the blot in (B) is plotted (data are means ± SEM of at least 3 independent experiments. ****p<0.0001 [Two-way ANOVA]). **E.** Assessment of the synchronized transport of the hGH-FM2-GFP in HeLa cells grown on indicated rigidities. Cells were treated as in (A) and the transport of the hGH-FM2-GFP visualised by video microscopy. Micrographs showing cells at the indicated trafficking time points. Dashed lines indicate the Golgi region. Scale bar, 20 μm. **F**. Quantification of the experiment in (E) shows the hGH-FM2-GFP fluorescence intensity in the Golgi region at different time points after D/D solubilizer and CHX addition. Data are means ± SEM.

HeLa-GH seeded on substrates with different stiffness (0.5 kPa, 8 kPa and >GPa) were exposed to ligand-induced cargo release from the ER in the presence of cycloheximide (CHX, 50 μg/ml). Western blot analysis revealed an enhanced secretion of hGH-FM2-GFP in cells grown on stiff substrate (>GPa), as shown by the detection of the reporter construct in the cell supernatant (Figure 2B, C). In contrast, the secretion of hGH-FM2-GFP was reduced in cells cultured on softer substrates, showing the most significant reduction observed on the 0.5 kPa compared to the 8 kPa substrate (Figure 2B, C), thus supporting the hypothesis of an intracellular trafficking delay. Importantly, upon measuring the intracellular levels of hGH-FM2-GFP, we obtained a similar result as previously observed for PC-I (Figure 1G-I). Cells grown on 0.5 kPa and 8 kPa displayed a significant decrease in the intracellular levels of hGH-FM2-GFP (i.e. 86% and 81%, respectively), after 2 hours of release from the ER (Figure 2B, D). This reduction was not accompanied by a corresponding increase in the protein levels of the cargo reporter in the media (Figure 2B, C).

In contrast, cells grown on rigid substrates (>GPa) exhibited a strong reduction in intracellular hGH-FM2-GFP levels, which reached approximately 96% after 2 hours (Figure 2B, D) and was accompanied, as expected, by the increase of the cargo reporter in the media (Figure 2B, C). Additionally, we analysed the secretion of TIMP1 and TIMP2 in HeLa cells using the Retention Using Selective Hooks (RUSH) system ^[42]^, which enables synchronized ER exit via biotin-induced cargo release. TIMP1- or TIMP2-eGFP RUSH constructs were retained in the ER through a streptavidin-KDEL hook and released upon biotin addition in the presence of cycloheximide (CHX, 50 μg/ml). Live-cell imaging confirmed synchronized trafficking of both cargoes (Figure S2A, B). We then seeded HeLa cells expressing TIMP1- or TIMP2-eGFP RUSH constructs on substrates with different stiffness (0.5 kPa, 8 kPa and >GPa) and induced cargo release with biotin. Western blot analysis revealed that, while intracellular cargo levels decreased under all conditions, efficient secretion into the medium occurred only on stiff substrates (>GPa). In contrast, secretion was impaired on softer matrices (0.5 and 8 kPa), confirming that substrate rigidity enhances trafficking and secretion efficiency across distinct cargoes and synchronization systems (Figure S2C-H).

The fact that the cargo reporters (i. e. hGH-FM2-GFP, TIMP1-eGFP and TIMP2-eGFP) are expressed under a constitutive promoter and the release of the cargo protein from the ER occurs in the presence of CHX (inhibitor of protein synthesis) argues against a possible transcriptional regulation induced by substrate stiffness. Instead, these observations suggest the involvement of a degradative pathway, in line with our data on PC-I (Figure 1G-I).

Given that matrix stiffness has been shown to increase intracellular tension ^[43,44]^ (see Figure 1A), regulating protein export from the ER through Rac1 ^[43,44]^, and also influences the final step of vesicular trafficking involving vesicle fusion with the PM ^[29]^, we set out to investigate whether Rac1 contributes to the stiffness-dependent regulation of ER export. To this end, HeLa-GH cells were treated with the Rac1 inhibitor NSC23766 and the trafficking of hGH-FM2-GFP from the ER was monitored using video microscopy. The inhibition of Rac1 delayed the arrival of hGH-FM2-GFP at the Golgi independently of the substrate rigidity on which the cells were grown (Figure S2I-L). This observation confirmed the hypothesis that substrate stiffness regulates protein transport from the ER to the Golgi in a Rac1 independent manner.

Based on these results, we focused then on the Golgi apparatus with the aim to investigate its role in stiffness-induced differential secretion. The specific involvement of the Golgi in this process remains unclear, except for a seminal study reporting that extracellular forces can influence Golgi rheology, and its response to applied forces ^[45]^. To address this, we seeded HeLa-GH cells on substrates of varying stiffness and track the intracellular trafficking of hGH-FM2-GFP by video microscopy. Our analysis revealed a significant delay in cargo transport in- and out the Golgi as substrate rigidity decreases (Figure 2E, F and Movie S1). We next investigated potential changes in Golgi morphology as a result of its substrates stiffness-controlled transport, revealing, however, no morphological changes by electron microscopy analysis (Figure S2M). Control experiments using the cis-Golgi marker GM130 and the trans-Golgi marker GalT1 showed no significant changes in protein localization or their expression levels, as determined by immunofluorescence (IF) (Figure S2N) and Western blotting (Figure S2O). Similarly, the protein levels of the ER-exit sites marker Sec31a and the Golgi proteins GRASP65 and GOLPH3 remained unchanged (Figure S2O).

Our findings indicate that substrate rigidity positively regulates the fraction of secreted proteins by suppressing their degradation.

### 2.3. Substrate stiffness regulates post-Golgi lysosomal degradation of secretory proteins

To test the involvement of degradation pathways in regulating the levels of secretory proteins during stiffness sensing, we first investigated the trafficking step at which cargo degradation occurs. To this end, we used a temperature block strategy to synchronise the trafficking of the hGH-FM2-GFP reporter. Specifically, the trafficking of the reporter was induced in cells incubated at 20 °C to accumulate cargo in the TGN ^[46]^, and then shifted to 37 °C to synchronously release cargo from the TGN (Figure S3A). Variation in substrate stiffness does not affect the intracellular levels of hGH-FM2-GFP when the cargo is blocked in the ER (Figure 3A-C, Sample 1) or during the trafficking of the cargo from the ER to the Golgi as confirmed by confocal microscopy and Western blot (Figure 3A-C, Sample 2). Instead, the intracellular levels of hGH-FM2-GFP drop during the transport out of the Golgi (Figure 3A-C, Sample 3). Specifically, hGH-FM2-GFP protein levels dropped by ∼75% 2 hours after resuming transport in cells grown on stiff substrate (>GPa), and by about 86% and 88% in cells grown on 8 kPa and 0.5 kPa substrates, respectively (Figure 3C). Under these conditions, the amount of hGH-FM2-GFP secreted in the media by cells grown on softer substrates was significantly reduced compared to the cargo secreted by those cells grown on a rigid substrate (Figure 3B). These results indicate that substrate stiffness controls hGH-FM2-GFP secretion by regulating post-Golgi cargo degradation. We also synchronized the trafficking of PC-I in BJ-5ta cells grown on substrates of different rigidities by applying a similar protocol (see Methods), using a temperature block at 20°C. We confirmed by confocal microscopy that the fluorescence intensity levels of PC-I dropped only during transport out of the Golgi, irrespective of the substrate on which the cells were grown (Figure S3D, E). This suggest, similarly to what was observed for hGH-FM2-GFP in HeLa-GH cells (Figure 3A-C), that the PC-I degradation occurring on soft substrates (see Figure 1G-I) takes place during post-Golgi transport, overall extending the generality of the mechanism to multiple cargoes and cell types. To evaluate the potential involvement of lysosomal degradation in this process, we assessed the trafficking and degradation of hGH-FM2-GFP in HeLa-GH cells seeded on substrates with increasing stiffness and treated with bafilomycin A1 (Baf A1), an inhibitor of lysosomal acidification ^[47]^. Western blotting data showed that Baf A1 treatment restored hGH-FM2-GFP protein levels in cells grown on 0.5 kPa and 8 kPa substrates, 2 hours after synchronized release from the ER (Figure 3D, E). In contrast, this recovery was not observed in cells grown on the rigid (>GPa) substrate. Imaging analyses further confirmed that Baf A1 caused accumulation of hGH-FM2-GFP in compartments positive for the lysosomal-associated membrane protein 1 (LAMP1) in cells grown on 0.5 kPa and 8 kPa substrates, whereas no such accumulation occurred in cells grown on the rigid substrate (Figure 3F, H). Notably, Baf A1 treatment induced a modest increase in hGH-FM2-GFP colocalization with the Golgi marker GM130 (Figure 3G, I), consistent with its known effect of impairing Golgi cargo export through disruption of organelle acidification ^[48]^. This effect was independent of substrate stiffness.

**Figure 3.**
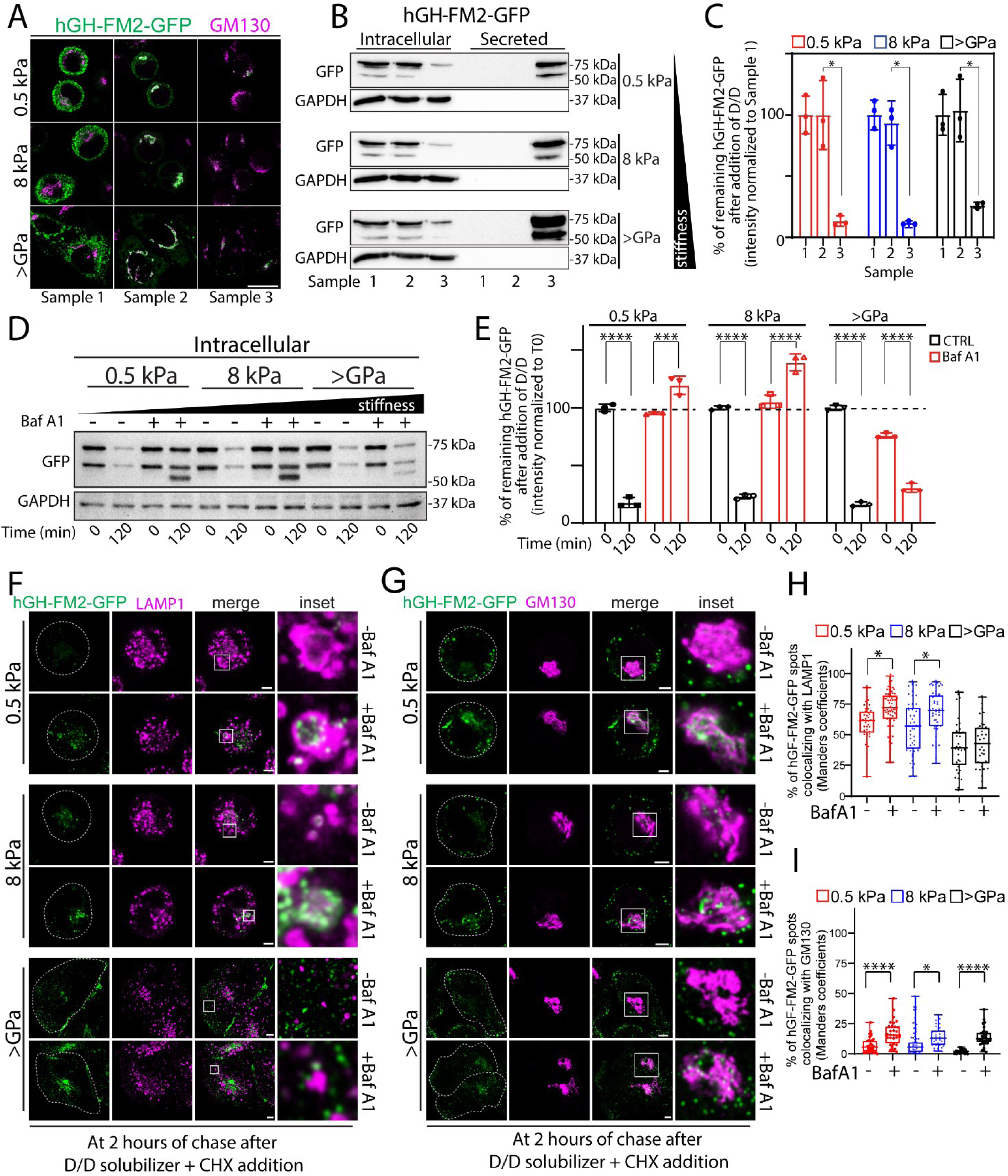
Substrate stiffness regulates post-Golgi lysosomal degradation of secretory proteins. **A**. HeLa-GH cells were seeded on indicated substrate rigidities and subjected to the synchronisation protocol described in (Figure S3A). Cells were then fixed, subjected IF and then imaged at each indicated time point and conditions (see Sample 1-3 in [Figure S3A]). hGH-FM2-GFP (green), GM130 (magenta). Scale bar, 20 μm. **B**. HeLa-GH were treated as in (A) and the cell lysate (intracellular) and the medium (secreted) were collected at indicated times and analysed by SDS-PAGE (20% of the total intracellular lysate and 100% of the total secreted was loaded, respectively) followed by Western blot (n = 3) with the indicated antibodies. **C**. Densitometric quantification of the blot in (B) is plotted (data are means of at least 3 experiments ± SD. *p<0.05 [Two-way ANOVA]). **D**. HeLa-GH cells were seeded on indicated substrate rigidities and treated with bafilomycin A1 (Baf A1, 20 n_M_) for 2 hours. Then, cells were exposed to D/D (1 μ_M_) and CHX (50 μg/ml) for 2 hours in the presence of Baf A1 (20 n_M_), lysed and processed for SDS-PAGE and Western blot with the indicated antibodies. Western blot shows intracellular levels of hGH-FM2-GFP at each indicated timepoint. Data are representative of at least 3 independent experiments. **E**. Densitometric quantification of the blot in (D) is plotted (data are means of at least 3 experiments ± SD. ***p<0.001, ****p<0.0001 [Two-way ANOVA]). **F, G.** HeLa-GH cells were seeded on indicated substrate rigidities and treated with Baf A1 (20 n_M_) for 2 hours. Then, cells were exposed to D/D (1 μ_M_) and CHX (50 μg/ml) for 2 hours in the presence of Baf A1 (20 n_M_), fixed and processed for IF labelling: hGH-GFP-FM (green) and LAMP1 (magenta) in (F); hGH-GFP-FM (green) and GM130 (magenta) in (G). Dashed lines indicate cells. White boxes indicate the inset. Scale bar, 5 μm. **H, I.** Quantification of hGH-FM2-GFP colocalization with LAMP1 lysosomal marker (H) from (F) or GM130 (I) from (G). Colocalization was calculated from >30 cells using ImageJ (JACoP plugin; Manders coefficients) (data are mean of at least 3 experiments ± SEM. *p<0.05, ****p<0.0001 [One-way ANOVA]).

We next investigated whether extracellular degradation might also contribute to the reduced secretion observed on soft substrates. We transferred conditioned medium containing secreted hGH-FM2-GFP, collected from cells grown on stiff substrates (>GPa) after 2 hours of trafficking activation, to cells cultured on 0.5 kPa, 8 kPa, and >GPa substrates. In these recipient cells, trafficking had also been previously activated for 2 hours. The medium was then replaced with the conditioned one and incubated for an additional 1 hour. Under these conditions, no degradation of hGH-FM2-GFP was observed, indicating that the reduced levels of the secreted protein on soft substrates are not due to extracellular degradation (Figure S3B, C).

These collective findings suggest that cells actively sense the stiffness of their microenvironment and respond to it by negatively regulating the trafficking of specific cargoes from the Golgi to the lysosomes, where proteins are degraded. Such traffic was proved to controls the flux and, therefore, the levels of proteins directed to the PM. These observations prompted us to dive into the mechanosensing pathway underlying this response.

### 2.4. Stiffness-driven driven changes in secretion and degradation are dependent on the Src-FAK axis

Focal adhesions (FAs) are specialised PM complexes enriched in integrin receptors that sense mechanical cues from the microenvironment and transduce them into a biochemical cellular response ^[49,50]^. Proto-oncogene tyrosine-protein kinase Src (Src) and focal adhesion kinase (FAK) are key effectors involved in FA-signalling ^[51–53]^, as they regulate stiffness-mediated mechano-transduction ^[51,54]^. Hence, we explored whether the effects of stiffness-driven changes in secretion were dependent on the tyrosine kinases Src and FAK. We first verified Src and FAK activation in HeLa-GH by assessing the stiffness-dependence of phosphorylation of FAK at Y397 and Src at Y418^[26,55]^ through Western blot analysis, and found this to be the case (Figure 4A-C). Specifically, both kinases exhibited robust activation on stiff substrates, while their activity was low on soft substrates (Figure 4A-C). Next, we investigated whether FAK and Src are necessary for the stiffness-induced changes in hGH-FM2-GFP secretion. To this end, we induced the synchronised transport of hGH-FM2-GFP in HeLa-GH silenced by RNA interference (RNAi) for Src (siSrc) or FAK (siFAK) and grown on substrates with different rigidities (Figure S4A). Then, applying the very same synchronization protocol, we monitored by Western blot, the intracellular and secreted hGH-FM2-GFP protein levels at time 0 and 2 hours after cargo release from the ER. We proved that both siSrc and siFAK cells grown on stiff substrate (>GPa) impaired the secretion of hGH-FM2-GFP compared to mock-treated cells, as confirmed by the reduced levels of the reporter construct in the media 2 hours after release from the ER (Figure 4D). To note, hGH-FM2-GFP was found to be highly accumulated intracellularly in siSrc cells, indicating a strong transport blockade. Conversely, siFAK cells completely lack intracellular hGH-FM2-GFP, which suggests its degradation (Figure 4D, E). Similar experiments performed in siSrc cells on 8 kPa substrate, revealed an intracellular accumulation of hGH-FM2-GFP and a modest block of its secretion 2 hours post-trafficking (Figure 4D, E). siFAK cells showed no notable effect under these conditions. Finally, the depletion of either Src or FAK kinase did not affect the intracellular or secreted levels of hGH-FM2-GFP on the softest substrate (0.5 kPa), compared to mock-treated cells, as expected (Figure 4D, E). These data indicate that Src and FAK silencing predominantly affects secretion in cells grown on rigid substrates, with minimal impact on softer substrates (0.5 kPa and 8 kPa). To further confirm these findings, we treated cells cultured on stiff substrates (>GPa) with either Dasatinib (Figure S4B-D) or FAK 14 (Figure S4E-G) inhibitors, which respectively block Src and FAK kinase activity ^[56,57]^. Strikingly, these pharmacological treatments phenocopied the effects observed upon RNAi-mediated depletion of Src and FAK, respectively (Figure S4B-G). This validates the specificity of our silencing approach and reinforces the conclusion that the kinase activities of Src and FAK are both stiffness-dependent and required for the regulation of cargo secretion and degradation.

**Figure 4.**
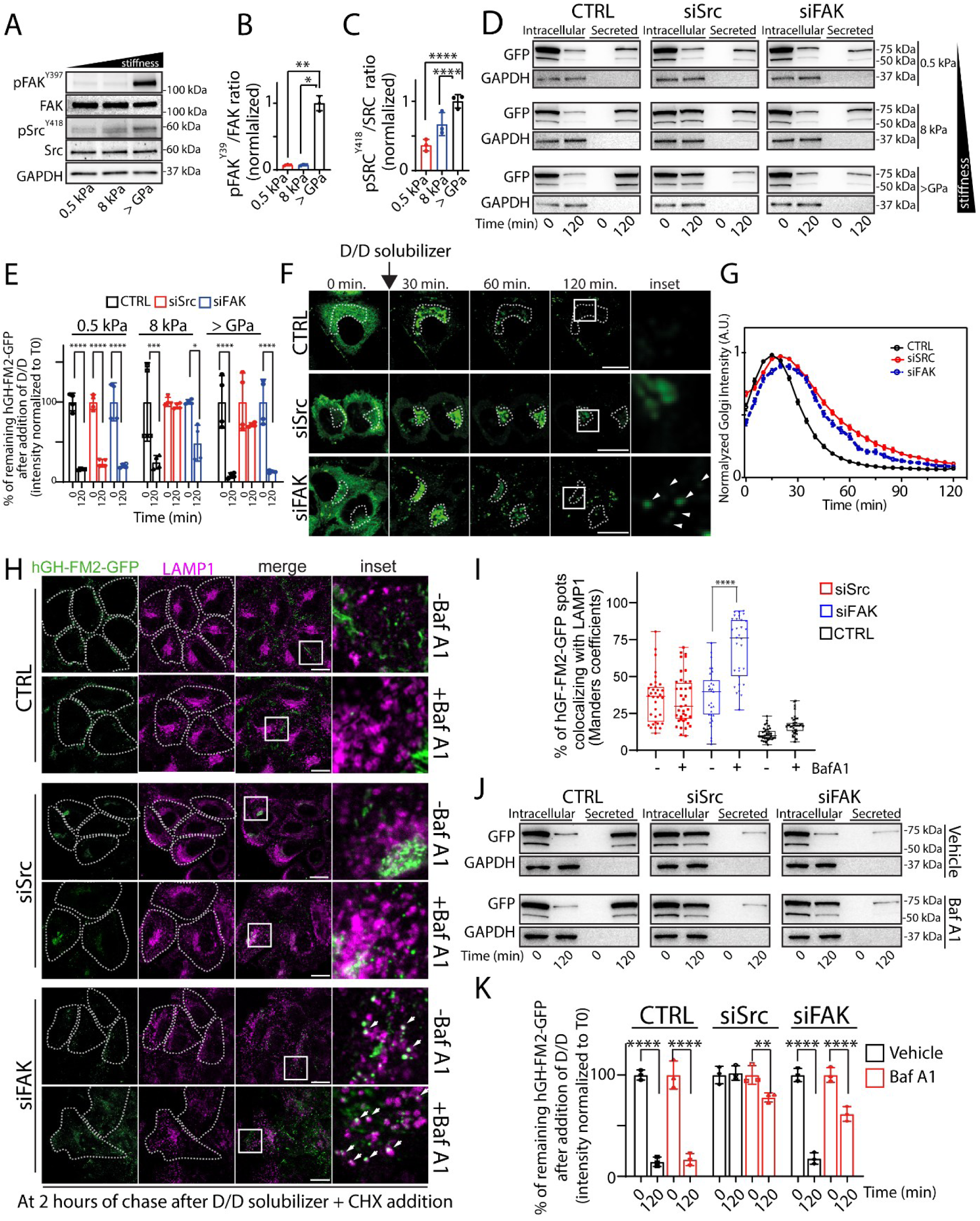
Stiffness-driven driven changes in secretion and degradation are dependent on the Src-FAK axis. **A.** Effects of substrate stiffness on FAK and Src. Cell lysates from HeLa-GH cells seeded on indicated substrate rigidities for 24 hours, then lysed and processed for SDS-PAGE and Western blotting with indicated antibodies. Data are representative of at least 3 independent experiments. **B, C**. The normalised ratio of p-FAK (Y397) to FAK (B) and of p-Src (Y418) to Src (C) in the experiment in (A), respectively (data are means of at least 2 experiments ± SD. *p<0.05, **p<0.01, ****p<0.0001 [One-way ANOVA]). **D.** HeLa-GH cells mock-treated (CTRL) or silenced for FAK (siFAK) or Src (siSrc) were grown on indicated substrate rigidities and then exposed to D/D (1 µM) and CHX (50µg/ml) for the indicated time points. The cell lysate (intracellular) and the medium (secreted) were collected at indicated timepoint and analysed by SDS-PAGE (90% of the total intracellular lysate and 100% of the total secreted was loaded, respectively) followed by Western blot with the indicated antibodies (n = 3). **E.** Densitometric quantification of the blot in (D) show the amount of intracellular hGH-FM2-GFP over time (data are means of at least 3 experiments ± SD. *p<0.05, ***p<0.001, ****p<0.0001 [Two-way ANOVA]). **F.** Assessment of the synchronized transport of the hGH-FM2-GFP in HeLa cells mock-treated (CTRL) or silenced for Src (siSrc) or FAK (siFAK) and grown on >GPa substrate. Cells were exposed to D/D (1 µM), CHX (50µg/ml) and then visualised by video microscopy. Micrographs showing cells at the indicated trafficking time points. White boxes indicate the inset. Arrowheads indicate hGH-FM2-GFP puncta. Dashed lines indicate the Golgi region. Scale bar, 20 μm. **G.** Quantification of the experiment in (F) shows the hGH-FM2-GFP fluorescence intensity in the Golgi region at different time points after D/D solubilizer and CHX addition. Intensity normalized to the maximum peak of the Golgi. Data are means ± SEM. **H.** HeLa-GH cells mock-treated (CTRL) or depleted of FAK (siFAK) or Src (siSrc) were grown on on >GPa substrate. After 24 hours cells were pre-treated with Baf A1 (20 n_M_) for 2 hours and then treated with D/D (1 µM) and CHX (50 µg/ml) for 2 hours in the presence of Baf A1 (20 n_M_), fixed and processed for IF labelling: hGH-FM2-GFP (green), LAMP1 (magenta). Dashed lines indicate cells. White boxes indicate the inset. Arrowheads indicate hGH-FM2-GFP and LAMP1 co-localisation. Scale bar, 20 μm. **I.** Quantification of hGH-FM2-GFP colocalization with LAMP1 lysosomal marker from (H). Colocalization was calculated from >31 cells using ImageJ (JACoP plugin; Manders coefficients) (data are mean of at least 3 experiments ± SEM. ****p<0.0001, [One-way ANOVA]). **J.** Cells were treated as in (H) and the cell lysate (intracellular) and the medium (secreted) were collected at indicated times and analysed by SDS-PAGE (90% of the total intracellular lysate and 100% of the total secreted was loaded, respectively) followed by Western blot with the indicated antibodies (n = 3). **K.** Densitometric quantification of the intracellular levels of hGH-FM2-GFP from the blot in (J) is plotted (data are means of at least 3 experiments ± SD. **p<0.01, ****p<0.0001 [Two-way ANOVA]).

We next performed time-lapse experiments in HeLa-GH grown on a stiff substrate (>GPa) to elucidate *i.* which step of hGH-FM2-GFP trafficking was blocked in siSrc cells, hindering its secretion and *ii*, the pathway through which the reporter was directed to degradation in siFAK cells. We found that hGH-FM2-GFP remained partly confined to the Golgi 2 hours after its release from the ER in both Src-depleted cells (Figure 4F, G and Figure S4H) and in cells treated with Dasatinib (Figure S4I, J), consistent with previous observations demonstrating the role of Src in regulating Golgi transport ^[58,59]^. Instead, FAK-depleted cells (Figure 4F, G and Figure S4H) and cells treated with FAK 14 (Figure S4I, J) displayed a cargo that was mildly confined within the Golgi and widely dispersed into cytosolic spots that were positive for the lysosomal marker LAMP1 (Figure 4H, I; Figure S4K). This dispersal was particularly pronounced in cells treated with Baf A1 (Figure 4H, I; Figure S4K). Western blot experiments were in line with these conclusions and confirmed that the inhibition of lysosomal degradation rescued the levels of hGH-FM2-GFP in siFAK cell, while no effects were observed in siSrc cells (Figure 4J, K). Finally, to elucidate the hierarchical relationship between Src and FAK in mechanotransduction, we performed Western blot analyses on stiff substrates (>GPa) using RNAi. The silencing of Src, either alone or in combination with siFAK, led to intracellular accumulation of hGH-FM2-GFP and impaired secretion (Figure S4L-N). These results position FAK downstream of Src in the pathway controlling stiffness-dependent cargo secretion and degradation.

Given the pivotal role of the Hippo signalling pathway in sensing and responding to mechanical cue of extracellular matrix (ECM) stiffness ^[60]^, we monitored the nuclear localization of Hippo-Yes-associated protein 1 (YAP) in HeLa-GH seeded on increased substrates with varying rigidity as a control experiment. As expected, the nuclear localization of YAP correlates with substrate stiffness (Figure S4O, P). However, pharmacological inhibition of actin polymerization and cytoskeletal contractility using cytochalasin D and Y-27632 treatments that are known to indirectly prevent YAP nuclear translocation (Figure S4Q, R), does not influence the secretion of hGH-FM2-GFP (Figure S4S, T). This suggests that the effect of matrix stiffness on hGH-FM2-GFP secretion is independent of YAP-mediated mechanotransduction.

Collectively, these experiments indicate that Src controls the export of hGH-FM2-GFP out of the Golgi, while FAK prevents hGH-FM2-GFP from being targeted to lysosomes for degradation, both in a stiffness-dependent manner.

These results confirm the existence of a refined cargo sorting control mechanism at the Golgi, driven by matrix stiffness and orchestrated by Src and FAK, which regulates the PM versus lysosomal trafficking of secretory proteins.

### 2.5. Golgi phospho-proteins differentially regulated during stiffness-sensing

To identify phospho-regulated mechanotransducers involved in cargo sorting at the Golgi downstream of Src and FAK, we employed an unbiased mass spectrometry (MS) phosphoproteomic approach (described in the Methods) to investigate proteome-wide changes in protein phosphorylation in HeLa-GH cells during stiffness-sensing. MS analysis identified 10,103 high-confidence class I phospho-sites across 1,600 phosphoproteins, with 58% phospho-enrichment (localization probabilities >0.75).

Principal Component Analysis (PCA) of the global phospho-proteomes revealed that cells cultured on soft substrates (0.5 kPa and 8 kPa) clustered closely with minimal separation along principal component 2 (PC2), whereas cells grown on stiff substrates (>GPa) formed a distinctly separate cluster along the same axis, suggesting a qualitatively different phospho-signature associated with high stiffness (Figure 5A; Dataset S1). Supporting this distinction, analysis of the overall phospho-site composition showed that serine (pS) residues were the most frequently phosphorylated, followed by threonine (pT) and tyrosine (pY). Phosphorylation levels were generally higher on soft substrates (except for pY, which was elevated under stiff conditions), indicating a stiffness-dependent shift in phosphorylation patterns (Figure S5A).

**Figure 5.**
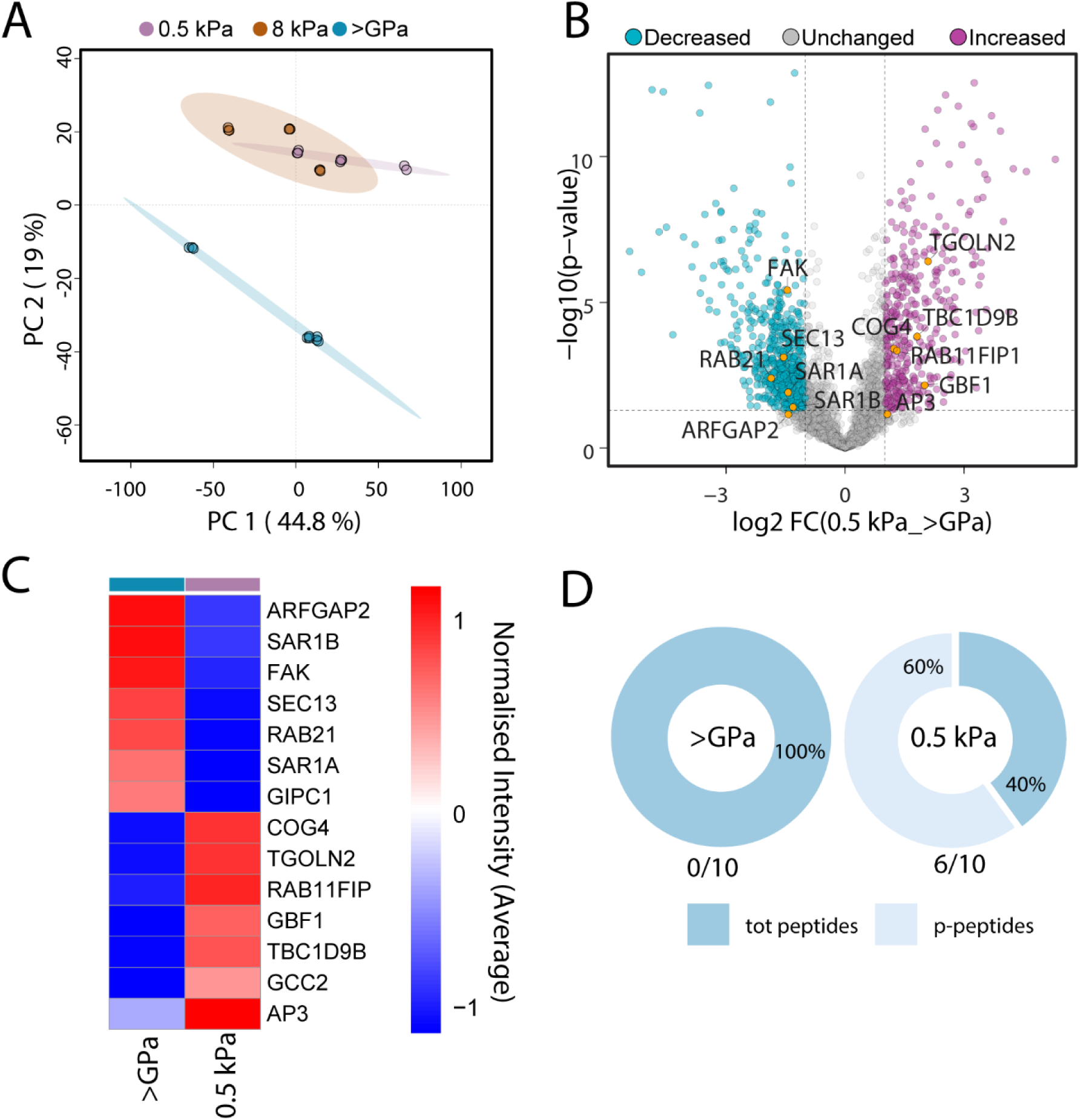
Effect of substrate stiffness on the phosphorylation dynamics of the Golgi proteome. **A.** PCA of the MS phospho-enriched proteomes from HeLa-GH cells grown on 0.5 kPa, 8 kPa and >GPa substrates, highlighting distinct quantitative abundance profiles. **B**. Volcano plot displaying differences in protein expression between 0.5 kPa and >GPa, with Golgi-related proteins highlighted in orange. **C.** Heatmap (Log2 LFQ normalized intensity) of the significantly deregulated phosphoproteins between 0.5 kPa and >GPa HeLa-GH cells. **D.** Pie charts illustrate the distribution of peptides associated with GBF1, separating phosphorylated from the total peptides (both phosphorylated and non-phosphorylated peptides).

To further investigate this observation, a comparative analysis across the three experimental groups was performed. Multiple testing identified 1,607 phosphoproteins as significantly deregulated (Figure S5B, Dataset S1). Cluster elaboration of the deregulated phosphoproteins revealed four distinct and biologically relevant quantitative trends (Figure S5B, Dataset S1). Gene Ontology Cellular Component (GOCC) analysis of proteins within each cluster indicated widespread deregulation of nuclear, cytoskeletal, and vesicular phospho-proteomes under soft substrate conditions (0.5 kPa and 8 kPa) compared to the stiff condition (>GPa), stress stiffness-dependent remodeling of key cellular compartments (Figure S5C, D). Notably, clusters 2 (upregulated) and 3 (downregulated) highlighted the key differences between stiff and soft conditions, while clusters 1 and 4 captured changes uniquely to the 0.5 kPa substrate (Figure S5C). Specifically, cluster 1 highlighted a broad upregulation of nuclear and cytoskeletal phosphoproteins in the 0.5 kPa condition, while cluster 4 revealed distinct subcellular localization patterns among downregulated phosphoproteins. Notably, phosphoproteins associated with the ER, lysosome, and Golgi were predominantly downregulated in cells cultured on 0.5 kPa substrate compared to both 8 kPa and stiff conditions (Figure S5D).

To further explore the functional implications of altered phosphorylation states across the three experimental groups, we examined the molecular activities associated with the deregulated proteins. GO Molecular Function (GOMF) investigation showed a generally balanced distribution of kinase and phosphatase activities across all clusters, except for cluster 4, which exhibited a marked enrichment of phosphatase activity (Figure S5E). This suggests that cells cultured on 8 kPa substrates maintain phosphorylation levels similar to the stiff control condition (>GPa), potentially due to active phosphatase regulation. In contrast, the 0.5 kPa condition stands out: here, phosphatase-associated proteins were downregulated, indicating a reduction in dephosphorylation activity. This likely contributes to the accumulation of phosphorylated proteins, consistent with the globally elevated phosphorylation observed in the phospho-enriched 0.5 kPa group (Figure S5A). These findings pointed to extensive remodelling of the phospho-proteome in response to substrate stiffness, with 0.5 kPa emerging as a critical softness threshold that triggers a distinct and consistent molecular shift. Based on this evidence, we focused our subsequent analyses on the 0.5 kPa and >GPa conditions to further dissect the underlying regulatory mechanisms. A direct comparison between these conditions revealed the deregulation of 968 phosphoproteins. Among these, we prioritized proteins known to be involved in the regulation of trafficking in and out of the Golgi apparatus. Proteins with increased phosphorylation on stiff substrates (>GPa) included GTP-binding protein Sar1A and Sar1B, Focal adhesion kinase 1 (FAK), PDZ domain-containing protein GIPC1 (GIPC1), RAB21, ADP-ribosylation factor GTPase-activating protein 2 (ARFGAP2), SEC13, while those with increased phosphorylation on soft substrates included Conserved oligomeric Golgi complex subunit 4 (COG4), trans-Golgi network integral membrane protein 2 (TGOLN2), TBC1 domain family member 9B (TBC1D9B), AP-3, GRIP and coiled-coil domain-containing protein 2 (GCC2), Rab11 family-interacting protein 1 (Rab11FIP1) and Golgi-specific brefeldin A-resistance guanine nucleotide exchange factor 1 (GBF1) (Figure 5B, C).

To validate the experimental setup, we inspected the well-known mechanosensitive phospho-site Y397 on FAK. Consistently, we observed highly significant phosphorylation in FAK Y397 on stiff substrate (>GPa) in both phospho-proteomes and Western blotting (Figure 4A, B).

Among the identified candidates, we prioritized coat proteins and their regulators known to be involved in cargo sorting. To assess their roles, we performed a plate reader-based secretion assay to measure the fluorescence intensity of hGH-FM2-GFP secreted into the media of cells grown on substrates of increasing stiffness (0.5 kPa 8 kPa and >GPa) and silenced for GBF1, Sar1b, SEC13, COG4, or GCC2 (see Methods; Figure S5F). Consistently, the secretion of fluorescent hGH-FM2-GFP in mock-treated cells positively correlated with substrate stiffness, confirming the reliability of the assay (Figure S5G). Silencing of Sar1b, SEC13, COG4, and GCC2 did not significantly affect hGH-FM2-GFP secretion in a clear substrate-dependent manner (Figure S5H). However, GBF1 silencing was highly detrimental to the cells, as previously reported, particularly in cells with high secretory activity ^[61]^. Despite this limitation, we observed a more pronounced inhibition of hGH-FM2-GFP secretion in cells grown on stiff substrates compared to those on soft substrates, suggesting a potential role for GBF1 in stiffness-dependent secretion (Figure S5H). Given the challenges associated with GBF1 silencing, we investigated the role of its phosphorylation.

### 2.6. GBF1 phosphorylation impacts the stiffness-dependent regulation of cargo sorting at the Golgi

GBF1 is a guanine nucleotide exchange factor (GEF) that, is essential for the formation of COPI vesicles at the early Golgi compartments, while at the TGN, it mediates BIG1- and BIG2-dependent Arf1 activation, necessary for recruiting clathrin adaptors ^[62,63]^. Inactivation of GBF1 by pharmacological (the fungal metabolite Brefeldin A, BFA) or genetic (siRNA against GBF1) manipulations results in the collapse of the Golgi into the ER and blocks secretion ^[60,61,64]^. Thus, GBF1 orchestrates cargo proteins selection and trafficking to their correct destination ^[64]^. Interestingly, recent studies have shown that Arf1, direct substrate of GBF1, exist in two functional different compartments involved in post-Golgi export of secretory cargo and endocytic recycling ^[59]^. GBF1 was notably responsive to substrate stiffness in HeLa-GH cells, showing significant upregulation on soft substrates (0.5 kPa) compared to rigid ones (> GPa). Of the peptides contributing to its identification and quantification, 60% were phosphorylated on soft substrates (0.5 kPa) but not on rigid ones (> GPa) (Figure 5D). Among these phosphorylated peptides, serine 1318 (S1318) was consistently phosphorylated in cells cultured on soft substrates (0.5 kPa) but absent in those grown on rigid substrates (> GPa) (Figure S5I). S1318 is the predominant phosphorylation site of GBF1, as reported in multiple studies (Figure S5J; https://www.phosphosite.org). Therefore, we hypothesised that GBF1 phosphorylation might impact the stiffness-dependent regulation of cargo sorting at the Golgi. To assess whether GBF1 phosphorylation on S1318 involved in the process of stiffness-dependent regulation of cargo sorting at the Golgi, we generated a phosphodeficient (S1318A) mutant of GBF1 and investigated its involvement in the stiffness-dependent regulation of protein secretion. To avoid potential effects of endogenous GBF1 in cells transfected with the wild-type GBF1 (GBF1-WT) or S1318A, we employed an assay to inactivate the endogenous GBF1^[64]^. To this end, both the GBF1-WT and S1318A constructs were also carrying the A795E mutation, which renders them resistant to BFA ^[64,65]^.

To get rid of the endogenous GBF1 activity, we first evaluated the ability of the two constructs to sustain Golgi architecture by exposing the cells to BFA. As shown by IF, only cells expressing the WT and S1318A constructs retained normal Golgi morphology, while cells expressing only the endogenous GBF1 shown a fragmented Golgi (Figure S6A, B).

To assess the trafficking capacity of cells expressing the GBF1 constructs, we performed the same GBF1 replacement assay in the presence of BFA and monitored the transport of hGH-FM2-GFP through the secretory pathway using live-cell video microscopy. The transport of the cargo reporter was observed exclusively in cells expressing either WT or S1318A constructs (Figure S6C and Movie S2, S3). In contrast, cells expressing only endogenous GBF1 (indicated with the asterisk in the Figure S6C and Movie S2, S3) showed a block in hGH-FM2-GFP trafficking, which remained entrapped in the ER as expected. These findings confirm the GBF1 replacement assay as a reliable method for studying the role of GBF1 phosphorylation in hGH-FM2-GFP secretion during stiffness sensing (0.5 kPa and >GPa). From this point onward, all experiments utilized BFA-resistant WT and mutant GBF1 under BFA treatment.

Next, we transfected HeLa-GH cells with GBF1-WT or S1318A mutant and we let them grow on substrates with different stiffness (0.5 kPa and >GPa) before inducing the trafficking of hGH-FM2-GFP. First, we confirmed by Western blot analyses that the expression levels of the two BFA-resistant constructs were similar (Figure S6D). During the transport assay, we observed that cells overexpressing the WT form of GBF1 and grown on soft substrate showed impaired secretion of hGH-FM2-GFP, when compared with that grown on a rigid substrate (Figure 6A, B). Interestingly, the replacement of endogenous GBF1 with the S1318A mutant in cells grown on soft substrates, restored the secretion of hGH-FM2-GFP to the levels of those observed in cells grown on stiff substrates (Figure 6A, B). This indicates that the GBF1 phosphorylation could regulate hGH-FM2-GFP degradation.

**Figure 6.**
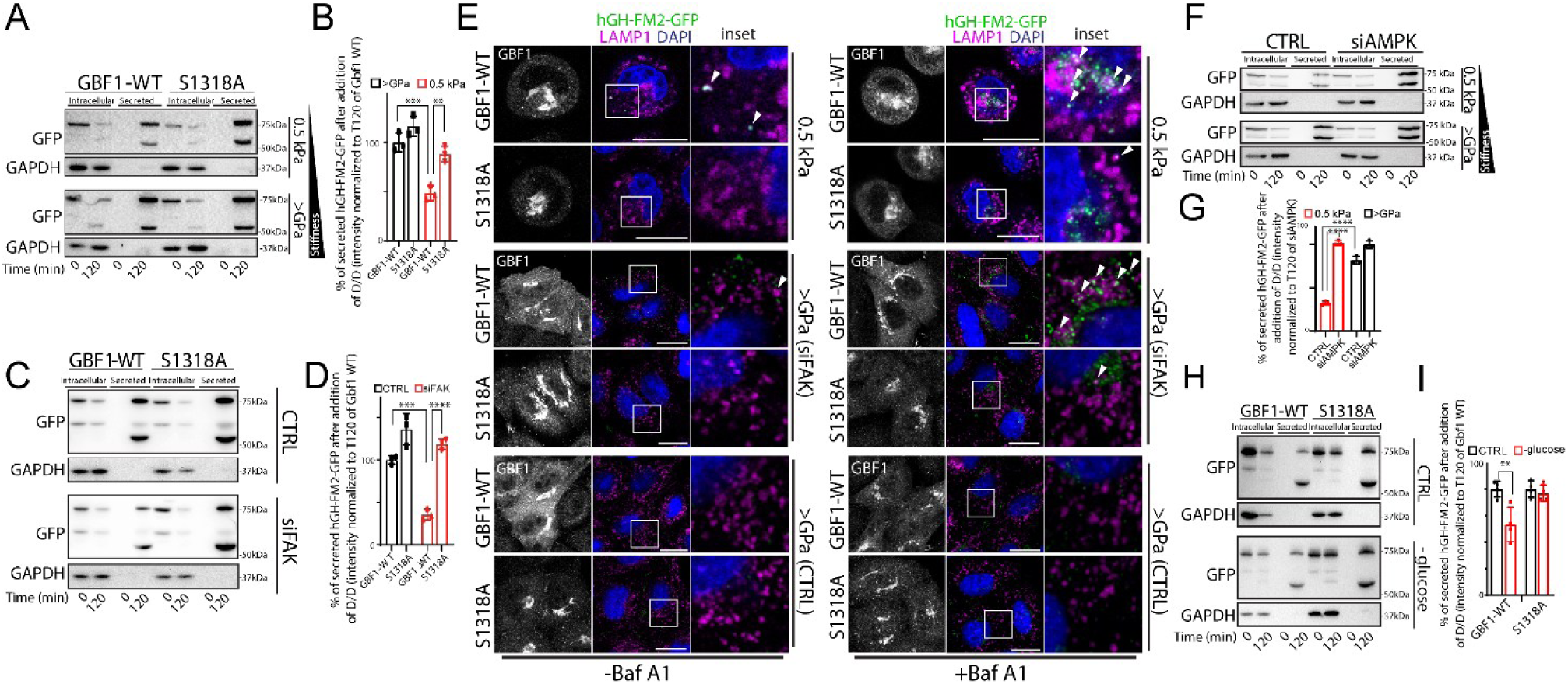
GBF1 phosphorylation impacts the stiffness-dependent regulation of cargo sorting at the Golgi. **A.** HeLa-GH cells expressing BFA-resistant GBF1-WT or S1318A mutant were grown on the indicated substrate. Cells were then treated with BFA (0.25 μg/ml) for 30 minutes prior to expose them to D/D (1 μ_M_), CHX (50 μg/ml) and BFA (0.25 μg/ml) for 2 hours. The cell lysate (intracellular) and the medium (secreted) were collected at indicated times and analysed by SDS-PAGE (20% of the total intracellular lysate and 100% of the total secreted was loaded, respectively) followed by Western blot with the indicated antibodies (n = 3). **B**. Densitometric quantification of (A) shows secreted hGH-FM2-GFP after 2 hours (data are mean of at least 3 experiments ± SD. **p<0.01, ***p<0.001 [Two-way ANOVA]). **C**. HeLa-GH cells mock-treated (CTRL) or silenced for FAK (siFAK) and expressing BFA-resistant GBF1-WT or S1318A mutant were grown on >GPa substrate. Cells were then treated with BFA (0.25 μg/ml) for 30 minutes prior to expose them to D/D (1 μ_M_), CHX (50 μg/ml) and BFA (0.25 μg/ml) for 2 hours. The cell lysate (intracellular) and the medium (secreted) were collected at indicated times and analysed by SDS-PAGE (20% of the total intracellular lysate and 100% of the total secreted was loaded, respectively) followed by Western blot with the indicated antibodies (n = 3). **D**. Densitometric quantification of (C) shows secreted hGH-FM2-GFP after 2 hours (data are mean of at least 3 experiments ± SD. ***p<0.001, ****p<0.0001 [Two-way ANOVA]). **E.** Mock-treated (CTRL) or siFAK HeLa-GH cells and expressing BFA-resistant GBF1-WT or S1318A mutant, were seeded on indicated substrates. Cells were treated with BFA (0.25 μg/ml) for 30 minutes prior to expose them to D/D (1 μ_M_), CHX (50 μg/ml) and BFA (0.25 μg/ml) for 2 hours, with or without Baf A1 (20 n_M_). Fixed cells were processed for IF labelling: hGH-FM2-GFP (green) and LAMP1 (magenta). White boxes indicate the inset. Arrowheads indicate colocalization. Scale bar, 20 μm. **F.** HeLa-GH cells mock-treated (CTRL) or silenced for AMPK (siAMPK) were grown on the indicated substrate. Cells were then treated with D/D (1 μ_M_) and CHX (50 μg/ml) for 2 hours. The cell lysate (intracellular) and the medium (secreted) were collected at indicated times and analysed by SDS-PAGE (20% of the total intracellular lysate and 100% of the total secreted was loaded, respectively) followed by Western blot with the indicated antibodies (n = 3). **G**. Densitometric quantification of (F) shows secreted hGH-FM2-GFP after 2 hours (data are mean of at least 3 experiments ± SD. ****p<0.0001 [Two-way ANOVA]). **H.** HeLa-GH cells expressing BFA-resistant GBF1-WT or S1318A mutant were grown on >GPa substrate (CTRL) and subjected to glucose starvation for 16 hours. Cells were then treated with BFA (0.25 μg/ml) for 30 minutes prior to expose them to D/D (1 μ_M_), CHX (50 μg/ml) and BFA (0.25 μg/ml) for 2 hours. The cell lysate (intracellular) and the medium (secreted) were collected at indicated times and analysed by SDS-PAGE (20% of the total intracellular lysate and 100% of the total secreted was loaded, respectively) followed by Western blot with the indicated antibodies (n = 3). **I.** Densitometric quantification of (H) showing the secreted hGH-FM2-GFP level (data are mean of at least 3 experiments ± SD. **p<0.01 [Two-way ANOVA]).

We then performed a similar experiment to test whether the S1318A mutant could restore hGH-FM2-GFP secretion in siFAK HeLa-GH on stiff substrates (>GPa), where the cargo reporter is directed towards the lysosomes (Figure 4H, I and Figure S4K), similarly to mock-treated cells on soft substrates (0.5 kPa). To assess this, we monitored hGH-FM2-GFP secretion in siFAK cells cultured on stiff substrates (>GPa) and transfected with either WT or S1318A mutants. Remarkably, the S1318A mutant restored hGH-FM2-GFP secretion levels to those of mock-treated cells on stiff substrate (>GPa), indicating a successful rescue of protein degradation (Figure 6C, D).

Furthermore, cells expressing GBF1-WT and cultured on soft substrates (0.5 kPa) showed hGH-FM2-GFP protein distributed within LAMP1 positive organelles at 2 hours post-trafficking (Figure 6E). By contrast, in cells expressing the S1318A mutant and grown on the same substrate, the cargo reporter was almost absent from LAMP1 compartments (Figure 6E) and was efficiently secreted with minimal intracellular retention (Figure 6A, B). Western blot experiments confirm that, Baf A1 treatment restored hGH-FM2-GFP protein levels in HeLa-GH expressing GBF1-WT and cultured on 0.5 kPa substrate 2 hours after synchronised release from the ER (Figure S6E, F). By contrast, this effect was absent in cells transfected with the S1318A mutant on the same substrate, as the mutant had already effectively prevented lysosomal degradation of the cargo reporter (Figure S6E, F). A similar rescue was seen in siFAK cells expressing the S1318A mutant on stiff substrates (>GPa) (Figure S6E, F), where the reporter was excluded from LAMP1 compartments and secreted, as shown by confocal imaging (Figure 6E). Together, these findings demonstrate that the GBF1 S1318A mutant overrides the degradation pathway, rerouting cargo to the PM for secretion, and position GBF1 downstream of FAK in the mechanotransduction cascade.

As a control, we investigated the role of GOPC in hGH-FM2-GFP trafficking during stiffness-sensing, as the phosphoproteomics analysis revealed an enhanced GOPC expression in cells cultured on >GPa substrates (PRoteomics IDEntifications database ID: PXD061714). This investigation was prompted by GOPC established role in post-Golgi sorting of cargos, such as syndecan-1 and CFTR ^[66,67]^. Although IF analysis of siGOPC HeLa-GH (Figure S7A) grown on a stiff substrate (>GPa) showed the mislocalization of hGH-FM2-GFP to lysosomes (Figure S7B), Western blot analysis demonstrated that GOPC silencing affects hGH-FM2-GFP secretion independently of substrate stiffness (Figure S7C, D). This highlights a specific role for GBF1 as a stiffness-dependent molecular switch for post-Golgi cargo sorting.

Taken together, these findings suggest that stiffness-dependent GBF1 phosphorylation at S1318 is critical for directing post-Golgi sorting either to the endo-lysosomal pathway for degradation or to the PM for secretion, thereby shaping the cellular secretome.

In searching for upstream GBF1 regulatory serine-threonine kinases, which are sensitive to variations in substrate stiffness, we first focused on candidates predicted to phosphorylate the S1318 residue of GBF1. Using prediction tools, we identified thirteen kinases with potential specificity for S1318, including AMPK, BARK1, CK1, CK2, CSNK2B, GRK, PHK, PRKCA, PRKCB, SIK1, SIK2, VRK1, and VRK2 (www.phosphosite.org). Heatmap analysis of our phosphoproteomic data revealed significant modulation of PRKCA, VRK1, CSNK2A1, CSNK2B, and PRKAA1 by substrate stiffness. Among these, AMPK (encoded by *PRKAA1*) caught our attention as it was already known to phosphorylate GBF1 at residue T1337 ^[68–70]^. AMPK was also known to be modulated by substrate stiffness and strongly activated in cells cultured on soft substrates ^[71]^. Consistently, we identified AMPK as one of the top phosphorylated proteins on soft substrates (0.5 kPa) (Figure S6G, H).

We investigated the role of AMPK inhibition in the synchronized transport of hGH-FM2-GFP in AMPK-silenced HeLa-GH cells (siAMPK; Figure S6I) cultured on substrates of varying rigidity (0.5 kPa or >GPa). Western blot analysis showed that siAMPK restored hGH-FM2-GFP secretion compared to mock-treated cells on soft substrates (0.5 kPa) (Figure 6F, G). Importantly, AMPK depletion had no effect on the secretion of hGH-FM2-GFP in cells grown on stiff substrates (>GPa) (Figure 6F, G). Conversely, AMPK activation in HeLa-GH cells on stiff substrates (>GPa), induced by glucose starvation or metformin treatment (Figure S6J), both well-established AMPK activators ^[72]^, significantly reduced hGH-FM2-GFP secretion, likely due to enhanced intracellular degradation (Figure S6K-M). Using the functional replacement assay in HeLa-GH grown on stiff substrates (>GPa) under AMPK-activating conditions, we observed that cells transfected with the GBF1-WT exhibited the expected reduction in hGH-FM2-GFP secretion (Figure 6H, I). In contrast, cells expressing the S1318A mutant restored cargo reporter secretion to comparable levels observed in cells transfected with GBF1-WT without AMPK activation (Figure 6H, I). This effect is due to the ability of the GBF1 mutant to bypass hGH-FM2-GFP sorting to LAMP1-positive compartments, despite AMPK activation (Figure S6N, O), placing GBF1 downstream of AMPK in the pathway.

To further support this model, we tested whether GBF1 and AMPK physically interact. Co-immunoprecipitation experiments performed in cells grown on soft substrates (0.5 kPa), where AMPK is active, demonstrated that AMPK co-precipitates with GBF1 (Figure S6P). Finally, to further clarify the signalling hierarchy, we silenced both AMPK and FAK in cells grown on >GPa substrate. While FAK silencing alone triggered hGH-FM2-GFP degradation, the combined knockdown restored its secretion (Figure S6Q, R).

Altogether these findings establish AMPK as a stiffness-dependent regulator downstream of FAK and upstream of GBF1 phosphorylation at S1318, necessary for the Gbf1-induced regulation of cargo lysosomal degradation and secretion. Activation of AMPK impaired hGH-FM2-GFP secretion, which can be restored by the S1318A mutant, highlighting the critical role of this phosphorylation site in connecting mechanotransduction with Golgi-sorting to regulate secretion.

## 3. Discussion

The secretory pathway synthesises and extracellularly releases thousands of proteins that mediate cell interactions with the environment and affect the physio-chemical properties of the environment itself. Despite all these information, how the cell senses and responds to biochemical and biophysical cues to regulate secretion remains a key question in cell physiology and pathology.

Here, for the first time we describe how substrate stiffness controls conventional protein secretion by regulating post-Golgi sorting through Src-FAK-AMPK-GBF1-dependent mechanosensing process, a mechanism relevant for numerous pathological conditions where the ECM or microenvironment are altered including fibrosis and solid tumours ^[73]^.

In response to increased matrix stiffness, we found an increase of nascent proteins and glycoproteins deposition on the PM. By using temperature-based or ligand-induced cell secretion assays, we found that substrate rigidity positively regulates secretion of several luminal cargo proteins, including collagen I, TIMP1-, TIMP2-eGFP RUSH and the cargo reporter hGH-FM2-GFP, in a transcriptional-independent manner. These dates align with previous findings indicating that matrix stiffness raises PM tension, which in turn enhances TIMP-1 exocytosis through dynamin-2 and caveolin-1^[29]^. Similarly, mechanical tension has been shown to regulate ER protein export through Rac1 ^[25]^, while ECM-related cues control COPII gene expression and ER-to-Golgi trafficking ^[74]^.

As such, the published data provide evidence for the regulation of protein trafficking by mechanical cues, with a focus on the early and late steps of secretion, which involve ER-exit and PM fusion of secretory carriers, respectively ^[29,74,75]^. However, while extracellular forces are known to impact Golgi rheology by reducing Arf1 recruitment ^[45]^, the specific role of the Golgi in this mechanosensitive context remains elusive. Here, we discovered a distinct mechanism that finely sense environmental mechanical cues and respond to stiffness variation by alternatively directing the secretory cargoes towards either degradation or secretion in an extremely precise fashion. The identification of GBF1 on TGN membranes ^[76]^, together with the recent discovery that its direct substrate Arf1 localizes to two functionally distinct TGN sub-compartments, further supports our findings. Both sub-compartments are decorated with clathrin, but differ in their clathrin-adaptor composition, which is key to directing cargo either to the PM or into the endocytic recycling pathway^[77]^.

Our imaging and biochemical experiments, performed by tracking the synchronised transport of the cargo reporter hGH-FM2-GFP and the physiological cargo PC-I, showed that substrate stiffness positively controls trafficking within and from the Golgi and negatively regulates the post-Golgi sorting to the lysosomes for degradation, thereby differentially affecting the secretion levels of specific cargoes. Our experiments demonstrate that Src and FAK, which are central regulators of cellular mechanosignalling ^[53,78,79]^, are activated in cells grown on a stiff substrate and to a lesser extent on softer substrates, in line with literature data ^[55,80]^. Combining pharmacological and genetic approaches with imaging and Western blot analysis, we found that Src inhibition or silencing impairs secretion on stiff substrates by disrupting protein export from the Golgi. In contrast, FAK inhibition or depletion promotes the rerouting of cargo proteins from the Golgi to lysosomes for degradation. On the contrary, the cells grown on soft substrate were much less affected by Src and FAK depletion. In searching for Golgi localized stiffness-induced regulators of protein secretion, we found that GBF1 is phosphorylated at Serine 1318 in response to decreased substrate stiffness using an unbiased phosphoproteomic approach. Overexpressing the S1318A GBF1 mutant in cells cultured on soft substrates or silenced for FAK hindered the sorting of hGH-FM2-GFP to lysosomes. In turn, this impairment increased the levels of the secreted cargo reporter, efficiently rescuing the secretion defect observed in cells grown on soft substrates or silenced for FAK and grown on stiff substrates.

It is noteworthy that S1318 of GBF1 falls within the consensus sequences recognized by several serine-threonine kinases (www.phosphosite.org). Among these, only AMPK, a master regulator of cellular metabolism ^[81]^, has been shown to directly phosphorylate GBF1, specifically at T1337, leading to Golgi fragmentation during mitosis ^[69,70]^. Moreover, AMPK has been localized to the Golgi, and its activation has been shown to inhibit the trafficking through the Golgi of vesicular stomatitis virus G (VSV-G) ^[70]^.

Our data demonstrates that AMPK interacts with GBF1 and its activation negatively regulate the secretion of hGH-FM2-GFP. This supports the hypothesis that matrix stiffness regulates GBF1-dependent secretion through AMPK, linking energy metabolism to mechanical stress and secretion. Consistent with our findings, the activation of GBF1 via AMPK has also been shown to influence the secretion and size of Weibel-Palade bodies in confluent monolayers of endothelial cells ^[68]^. AMPK was reported to be activated in cells cultured on soft ECM stiffness ^[71]^ and inhibited on stiff substrates through Src-FAK-mediated phosphorylation ^[82]^. Our data support a model in which increasing substrate stiffness triggers a positive feedback loop. In this mechanism, the activation of Src and FAK kinases suppresses AMPK-dependent phosphorylation of GBF1, thereby reducing the lysosomal targeting of ECM components and ECM-remodelling factors (Figure S1F-H) and promoting their secretion instead (Figure 7A, B). As a result, a vicious cycle is established: increasing ECM stiffness promotes augmented secretion of its components, which further contributes to the stiffening of the ECM. Mechanistically, the data align with a more complex picture in which GBF1 phosphorylation at S1318 may trigger the ARF1 dissociation from the Golgi ^[45]^, as observed in previous studies when cells are cultured on soft substrates. This, in turn, could redirect cargo sorting toward lysosomes for degradation ^[83]^. Although we identified S1318 as a key mechanosensitive site, GBF1 is phosphorylated at many other residues whose roles remain unclear ^[65]^. Other sites, such as T1337, linked to membrane dissociation, or those affecting GBF1 stability and function, may also regulate Golgi export or compartment-specific Arf1 activation ^[70]^. Additionally, GBF1 localization within Golgi subdomains may cooperate with its phosphorylation state to control cargo fate under mechanical stress. Further studies are needed to understand how these factors integrate to regulate post-Golgi trafficking.

**Figure 7.**
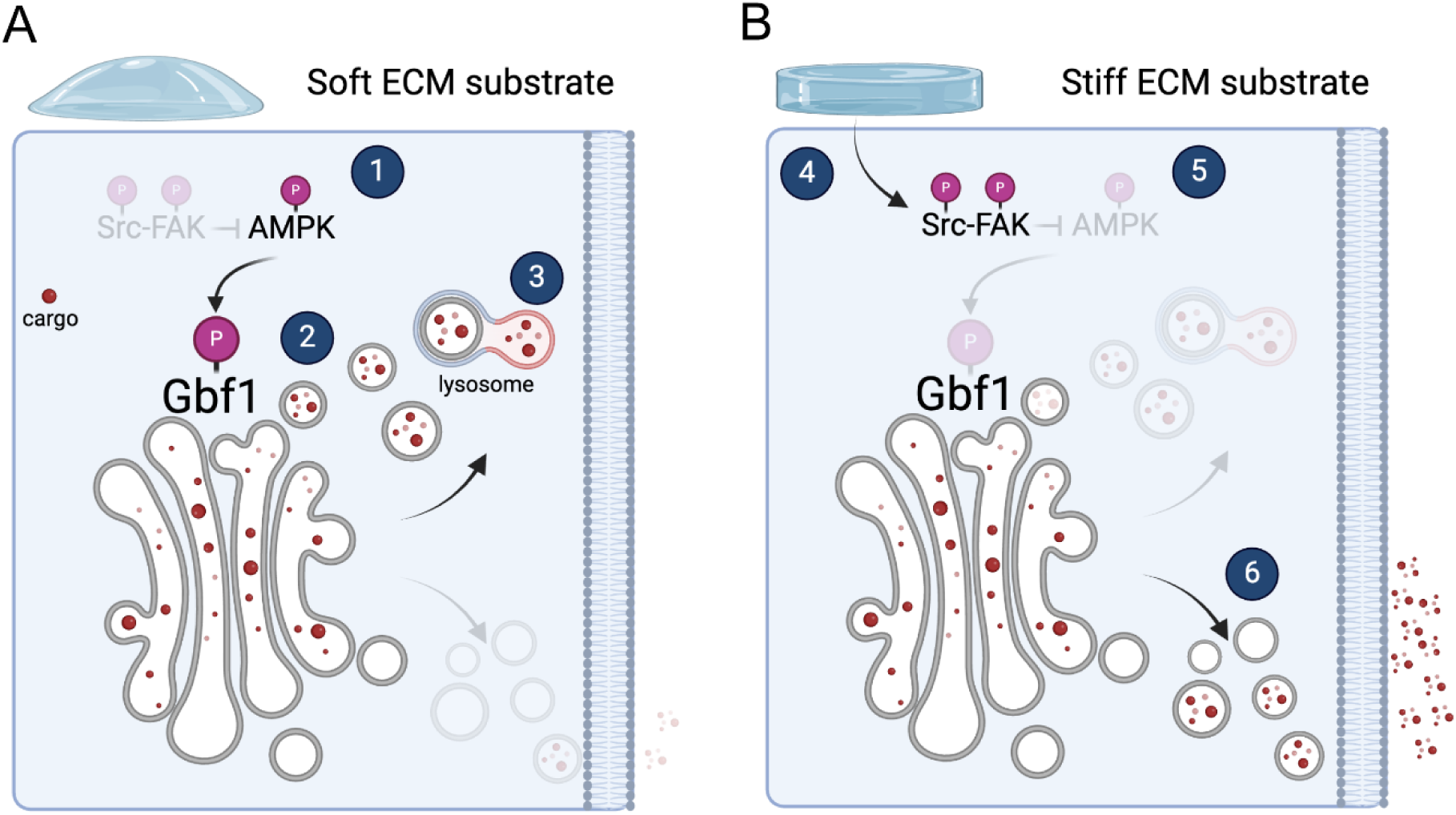
Proposed model showing regulation of secretion via the Src-FAK-GBF1 axis in response to ECM stiffness. **A**. On soft matrices, the inactivity of Src and FAK kinases (1) permits AMPK phosphorylation at T172, which may, in turn, phosphorylate GBF1 at S1318 (2). Phosphorylated GBF1 directs cargoes toward lysosomal degradation rather than secretion (3), resulting in a significant reduction in secretion levels. **B**. In contrast, on stiff matrices, mechanosensors activate Src and FAK kinases (4), inhibiting AMPK phosphorylation at T172 (5). The resulting decrease in AMPK activity may prevent GBF1 phosphorylation at S1318, blocking cargo sorting to lysosomes. Consequently, cargoes are secreted rather than degraded (6), (created with BioRender.com).

Additional regulators could also participate in this mechanosensitive pathway, with mTOR representing an intriguing candidate given its dual role in mechanosensing and trafficking regulation. Since mTORC1 is activated by mechanical tension via FAK-PI3K-AKT ^[84,85]^ and inhibited by AMPK ^[86–88]^, mTOR could function as a molecular switch integrating these opposing signals to control cargo fate at the Golgi. We speculate that on stiff substrates, mTORC1 activation might promote secretory trafficking by counteracting AMPK-mediated GBF1 phosphorylation, while mTOR inhibition on soft substrates could favour lysosomal targeting of specific secretory cargoes. This model is supported by our observation that AMPK knockdown rescues secretion in cells grown on soft substrates, suggesting an antagonistic AMPK-mTOR axis controlling post-Golgi sorting decisions.

The identification of this Src-FAK-AMPK-GBF1 axis reveals a sophisticated post-translational control mechanism that allows cells to rapidly adapt their secretory activity in response to environmental changes. This regulation operates within minutes to hours, potentially preceding slower, transcription-dependent responses. Physiologically, this regulation could be utilized by cells experiencing rapid changes in stiffness, such as immune cells like platelets, neutrophils, and dendritic cells to enhance secretion of proteins such as metalloproteinases during tissue infiltration ^[89,90]^. Conversely, in pathological conditions like fibrosis, this acute response becomes maladaptive, as increased matrix stiffness promotes excessive ECM secretion, further increasing tissue stiffening. The components of this circuit have already been found to be dysregulated in several pathologies characterized by impaired secretion. AMPK activation has been reported to play a protective role in fibrosis^[91–94]^. Recently, a rare genetic variant of GBF1 was identified in certain families with early-onset familial pulmonary fibrosis, highlighting the potential relevance of this gene in lung fibrosis ^[95]^. Therefore, investigating the interplay between AMPK and GBF1 in the context of wound healing and fibrosis would be particularly intriguing. The existence of a feedback loop between ECM stiffness and cell-mediated deposition and degradation of matrix proteins is well established, particularly in terms of transcriptional regulation and activity. One example is represented by the transcriptional regulator NFATC4, that has been shown to regulate the stiffness dependent myofibroblast collagen secretion ^[96–99]^. However, there is still limited understanding of how secretory pathway regulation fits into this framework ^[96]^. Phosphorylation of GBF1 could represent a promising target for disrupting the positive feedback loop between matrix stiffness and aberrant secretion, a hallmark of fibrotic diseases ^[100,101].^

From a broader perspective, our findings offer novel insights into the general mechanism of secretion, going beyond ECM-related cargoes. Specifically, we reveal a novel layer of post-translational regulation that influences the sorting of selected cargoes at the TGN, impacting the overall cell secretome. These discoveries raise several important questions: i. Which cargoes are sensitive to this regulation? ii. Do these cargoes have a recognition signal for sorting? iii. What are the trafficking components at the TGN that recognize these sorting signals? iv. How is this recognition achieved for luminal cargoes?

Moreover, this study expands our understanding of the pathological mechanisms involved in dysregulated ECM deposition, which can lead to excessive wound healing and fibrosis, ultimately affecting tissue function.

## 4. Methods

### Reagents and plasmids

Cycloheximide was from ThermoFisher Scientific, D/D Solubilizer was from Clontech Laboratories Inc. Cycloheximide, Brefeldin A, Bafilomycin A1, Metformin, Cytochalasin D, Y-27632, NSC23766, Fibronectin, Dasatinib, FAK 14 inhibitor, L-azidohomoalanine (AHA), N-azidoacetylmannosamine (ManNAz), and Dibenzocyclooctyne-Cy5 (DBCO-Cy5) were from Sigma Aldrich. Paraformaldehyde 8% Aqueous Solution, EM Grade was from Electron Microscopy Science. T-75 flasks, multiwells and coverslips with 0.5 kPa and 8 kPa elastic modules were from CytoSoft, Advanced BioMatrix. Substrates with stiffness indicated as >GPa were either glass coverslips or plastic flasks or multiwells.

The GBF1 cDNA used in this study has been described previously ^[102]^. GBF1 A795E, from ^[76]^, was cloned in the ptd-Tomato N1 (Clontech), and GBF1 (S1318A) mutation was introduced using the Phusion Site-Directed Mutagenesis Kit (F541) from the Thermo scientific. The substitution was confirmed by sequencing. The sequences of the primers used for site-directed mutagenesis were: S1318A forward (5’-GTACACTGCCGACTCAGAGGTCTACACTGACCATGGCAGG-3’) S1318A reverse (5’-GAGTCGGCAGTGTACCCTCGATCCAGGCTCACGTCATTCT-3’).

The open reading frames (ORFs) of TIMP1 (CloneID OHu24051; NM_003254.3) and TIMP2 (CloneID OHu16561; NM_003255.5) were obtained from Genscript and subcloned by the provider into the RUSH plasmid backbone, upstream of eGFP.

### Cell lines and culture conditions

HeLa-M (human cervical cancer cells, female origin) were obtained from the ATCC and grown in RPMI-1640 supplemented with 10% FCS. HeLa-M cells stably expressing hGH-FM2-GFP (HeLa-GH) were from our previous studies ^[18]^. HeLa-GH were grown in RPMI (Gibco, USA) supplemented with 10% fetal calf serum (FCS) (Gibco). BJ-5ta (human immortalized fibroblasts, male origin) were obtained from the ATCC and grown in DMEM: Nutrient Mixture F-12 (DMEM F-12) supplemented with 10% FCS. All media were supplemented with penicillin-streptomycin and L-glutamine. Cells were cultivated at 37°C and 5 % CO2 in a humidified atmosphere. Before seeding, the plates were coated with fibronectin (10 μg/ml) for 30 minutes to help the cells stick to the different substrates with different elastic modules.

### Nascent glycoproteins labelling

The protocol has been previously used by ^[103]^. Briefly, HeLa cells were seeded on fibronectin-coated coverslips with elastic modules, 0.5, 8 kPa or >GPa cultured in Dulbecco’s Modified Eagle Medium (DMEM; Gibco) supplemented with 10% fetal bovine serum (FBS) and 1% penicillin-streptomycin at 37 °C in a 5% CO₂ incubator. Upon reaching ∼70-80% confluency, the cells were washed twice with pre-warmed phosphate-buffered saline (PBS) to remove residual methionine from the standard growth media. To replace methionine and enable incorporation of L-azidohomoalanine (AHA) cells were incubated with methionine-free DMEM supplemented with 10% dialyzed FBS (to eliminate methionine) and 50 µM AHA (Thermo Fisher Scientific). After 24 hours of labelling of nascent proteins, cells were fixed in 4% paraformaldehyde (PFA) in PBS for 15 minutes at room temperature. Similarly, to measure the synthesis of nascent glycoproteins, HeLa were grown in presence of 50 µM N-azidoacetylmannosamine (ManNAz) for 24 hours. The AHA-or ManAz-labelled proteins were then conjugated with Dibenzocyclooctyne-Cy5 (DBCO-Cy5, 10 µM) in PBS with 1% BSA for 30 minutes at room temperature, protected from light to avoid photobleaching. After staining, excess DBCO-Cy5 was removed by washing the coverslips three times with PBS. Nuclear staining to visualize nuclei, cells were incubated with DAPI (Thermo Fisher, 5 µg/ml in PBS) for 10 minutes at room temperature. Excess dye was washed off with PBS immediately prior to imaging.

### Cytokine secretion

Synchronisation protocol for protein secretion: HeLa cells were seeded on silicone substrates with elastic modulus of 0.5 kPa, 8 kPa, or >GPa for 24 hours and then incubated for 3 hours at 10 °C to achieve a general the block of the trafficking and accumulation of the cargoes within early compartments (ER and ERGIC) of secretory pathway ^[104,105]^. Then the temperature was shifted to 37°C for 2 hours in the presence of cycloheximide (CHX, 50 μg/ml) to achieve a synchronized secretion of cargos. After 2 hours of chase, the medium was collected and used for cytokine array analysis (Ray Biotech, Human Cytokine Array C5) according to the manufacturer instructions.

### Synchronization protocols

After HeLa-GH were grown for 24 hours on the specified substrate stiffness. To achieve simultaneous hGH-FM2-GFP cargo release from the ER, D/D Solubilizer (1 μ_M_) was added to the cells for the durations shown in each figure ^[40]^, in the presence of cycloheximide (CHX, 50 μg/ml). Then the cells were fixed and subjected to immunofluorescence or alternatively lysed and subjected to SDS-PAGE and Western blotting.

Alternatively, HeLa cells were transfected for 24 hours with TIMP1 or TIMP2-eGFP-RUSH plasmids and then grown for 24 hours on the specified substrate stiffness. To achieve simultaneous cargo release from the ER, biotin (80 μ_M_) was added to the cells for the durations shown in each figure ^[42]^, in the presence of cycloheximide (CHX, 50 μg/ml). Then the cells were fixed and subjected to IF or alternatively lysed and subjected to SDS-PAGE and Western blotting.

For the traffic assay to synchronised ER-to-Golgi or Golgi-to-PM transport of hGH-FM2-GFP cargo, HeLa-GH were grown for 24 hours on the specified substrate stiffness (Sample 1 in Figure 3A). Alternatively, D/D solubilizer (1 μ_M_) was added to the cells for 2 hours at 20 °C to achieve accumulation of the cargo to Trans-Golgi Network (TGN) (Sample 2 in Figure 3A), or to cells treated with D/D solubilizer (1μ_M_) for 2 hours at 20 °C were kept for further 2 hours at 37 °C to achieve the release of the cargo from the TGN and trafficking to PM (Sample 3 in Figure 3A). After each trafficking protocol the cells were fixed and subjected to immunofluorescence or lysed and subjected to SDS-PAGE and Western blotting.

Synchronous Col1A1 release from the ER of BJ-5ta fibroblast cultured on indicated substrate stiffness for 24 hours was performed by washing the cells 3 times with PBS and keeping them for 3 hours at 40 °C in DMEM-HEPES supplemented with 1% FCS without ascorbic acid. A traffic pulse (60 minutes) of Col1A1 was induced by shifting cells to 32 °C in the presence of 100 μg/ml ascorbic acid ^[106]^. Alternatively, for synchronization of PC-I trafficking from the Golgi to the PM, after the 3-hour incubation at 40 °C, cells were shifted to 20 °C for 2 hours to block Golgi export, followed by incubation at 32 °C for an additional 2 hours to allow trafficking to resume. Following to these treatments, cells were subjected to SDS-PAGE and Western blotting. Cycloheximide (CHX; 50 μg/ml) was added 30 minutes before all traffic pulse experiments to ensure that most of the secretory cargo monitored and the response it generates in the cell was specific.

### Western blot and Immunoblotting

For Western blot analysis, cells were lysed in RIPA buffer (50 mM Tris-HCl pH 7.6, 150 mM NaCl, 1 % NP-40, 0.1 % SDS, 0.5 % sodium deoxycholate) supplemented with complete protease inhibitor (Roche Diagnostics, Basel, Switzerland) and phosphatase inhibitor (PhosStop, Roche Diagnostics), incubated on ice for 30 minutes and centrifuged at 14,000 g for 10 minutes at 4 °C. Protein concentrations were determined using the PierceTM BCA Protein Assay Kit (ThermoFisher Scientific). 6x Laemmli loading buffer (6% SDS, 300 mM Tris-HCl, pH 6.8, 25 % (w/v) glycerol, 5% β-mercaptoethanol, 710 mM, 0.1% bromophenol blue containing 50 mM NaF and 100 mM ß-glycerol phosphate) was added to samples (10-20 μg protein lysate) and heated at 95 °C for 5 minutes. Lysates were loaded on 10-15% SDS-PAGE gels and blotted onto nitrocellulose membranes. Electrophoretic separation was performed at 100-120 V, RT and transfer to nitrocellulose membranes was performed at 350 mA for 2 hours on ice. Membranes were blocked and incubated with primary antibodies (see Table S1) overnight at 4°C and subsequently with secondary antibodies 1 hour at room temperature (1:20,000, Santa Cruz). ChemiDoc imaging system (BioRad Laboratories, Inc.) was used to detect chemiluminescence. Densitometric quantifications were performed by using ImageJ software.

### Co-immunoprecipitation

For co-immunoprecipitation experiments, cells overexpressing GBF1-WT and grown on soft 0.5 kPa substrate were crosslinked using dithiobis(succinimidyl propionate) (DSP; Thermo Fisher) freshly prepared at a stock concentration of 125 mg/ml in DMSO. For crosslinking, DSP was diluted 1:125 in pre-warmed PBS to obtain the working solution. Then, cells were washed once with PBS before adding the DSP solution and incubated at room temperature (RT) for 7 minutes. To quench the crosslinking reaction, Tris-HCl (pH 8.5) was added directly to the culture dish, followed by a 1-minute incubation at RT. Cells were then washed twice with PBS and lysed directly in ice-cold RIPA buffer (150 mM NaCl, 1% NP-40, 0.5% sodium deoxycholate, 0.1% SDS, 1 mM EDTA, 50 mM Tris HCl pH 8) supplemented with protease and phosphatase inhibitors. Total lysates were passed ten times through a 25-gauge needle with syringe, kept at 4 °C for 30 min, processed with three freeze-thaw cycles, and then cleared by centrifugation in a microcentrifuge (18,800 g at 4 °C for 20 min). The lysate was cleared and quantitated using the BCA Protein Assay kit (Pierce).

For immunoprecipitation, 1.5–3 mg of total protein were incubated with 1-3 µg of anti-GBF1 antibody (see Table S1) over night at 4°C with gentle rotation, then incubated for 2 hours at 4 °C with ProteinG-Sepharose. Then, the samples were washed extensively with immunoprecipitation buffer and analysed by SDS–PAGE and immunoblotting with anti-GBF1 and anti-AMPK antibodies (see Table S1).

### siRNA treatments and transfections

The siRNAs utilized in this study were obtained from Sigma-Aldrich. Oligofectamine (Invitrogen) was used for siRNA transfection and performed according to manufacturer’s instructions (used at concentrations between 5-50 n_M_ for the target gene for 72 hours). Each individual siRNA in the pool was also tested in localization and signaling experiments. The list of siRNA sequences can be found in Table S2. The silencing efficiencies were evaluated either by Western blotting or by qPCR using specific primers (see Table S3). HeLa-M cells stably expressing hGH-FM2-GFP were transfected with BFA-resistant GBF1 constructs with Mirus or Lipofectamine® LTX transfection reagents following the manufacturer’s instructions.

### RNA extraction and real-time qPCR

Total RNA was isolated from cells using RNeasy Mini kits (Qiagen) following the manufacturer’s instructions. The quality and quantity of RNA was determined by NanoDrop 2000c spectrophotometer (Thermo Scientific) and TAE agarose gel electrophoresis, respectively. Total RNA was reverse transcribed to cDNA using the QuantiTect Reverse Transcription Kit (Qiagen) and subjected to real-time qPCR using specific primers for the target genes of interest in the presence of LightCycler® 480 SYBR Green I Master Mix (Roche) on a LightCycler® 480 II detection system (Roche). Hypoxanthine phosphoribosyltransferase 1 (HPRT1) was used as the reference gene and the variations in the expression levels of target genes were determined and expressed as fold changes respect to the control, using the comparative CT method (ΔΔCT method). The list and sequences primers used in this study can be found in Table S3.

### Electron microscopy

The sample preparation was done essentially as described previously ^[107]^. In brief, HeLa cells were seeded on indicated substrate rigidities for 24h, fixed with with 4% paraformaldehyde and 0.05% gluteraldehyde, for 30 minutes at room temperature, and the samples were then dehydrated and epon embedded. The samples were sectioned (200-nm sections) and examined by an electron microscope (Tecnai-12; FEI). Random sampling for images showing different the compartments of the secretory pathway was performed. Images were identified and acquired.

### Phosphoproteome sample preparation

Phosphoproteomic analysis was conducted using four independent biological replicates, each prepared on separate days by seeding cells onto substrates with different stiffness levels. CytoSoft T-75 flasks (Advanced BioMatrix) with elastic moduli of 0.5 kPa, 8 kPa, or standard tissue culture plastic were coated with 10 μg/ml fibronectin for 1 hour at room temperature. A total of 5 × 10⁶ cells were seeded per flask and incubated overnight. Cells were then lysed in 1 ml of 8 M urea, 50 mM Tris-HCl supplemented with complete protease and phosphatase inhibitors. Phosphopeptide enrichment was performed using about 2 mg of total proteins from each sample lysate, to which 50 mM ammonium bicarbonate (Ambic) was added to produce a final volume of 1 ml. Thus, the samples were subjected to in-solution digestion and enrichment in phosphopeptides according to the following solution digestion and desalting process. Each sample received 50 μL of 200 mM tris(2-carboxyethylphosphine) (TCEP), which was added and incubated for an hour at 55°C. Next, 50 μL of 375 mM iodoacetamide (IAA) was added, and the sample was left to incubate for 30 minutes at room temperature in the dark. 50 μg of trypsin (enzyme:protein ratio 1:40) was added and each sample and then incubated at 37°C overnight with shaking (600 rpm). 10 μL of 100% trifluoroacetic acid (TFA) were added, and the concentration of the samples was performed, until completely dry, using a speedvac. To the samples were added 300 μL of TFA 0.1% and then subjected to a desalting procedure using Pierce™ Peptide Desalting Spin Columns (cat no. 89852) following the manufacturer’s instructions. The eluates were concentrated by speedvac until they were completely dry. Each biological replicate was then divided into four technical replicates prior to phosphopeptide enrichment.

### Phosphopeptides enrichment

The samples were subsequently enriched in phosphopeptides according to the following procedure. Samples were resuspended by addition of 150 μL of Binding-Equilibration buffer, and pH was corrected to pH < 3. Phosphopeptide enrichment was performed with High-Select™ TiO2 Phosphopeptide Enrichment Kit (cat no. A32993) according to the manufacturer instructions.

The eluates were concentrated by speedvac until they were completely dry and then resuspended in of 40 μL of 0.1% formic acid (FA). Protein quantification was performed by using the Pierce™ Quantitative Colorimetric Peptide Assay Kit. The samples were dried again by speedvac and resuspended in 0.1% FA to obtain samples with a peptide concentration equal to 0.5 μg/μL. The samples prepared in this way were analyzed by nanoLC-HRMS under the conditions reported below.

### High resolution mass spectrometry analysis (nLC-HRMS)

All samples were analyzed at UNITECH OMICs (University of Milano, Italy) using a Dionex Ultimate 3000 nano-LC system (Sunnyvale, CA, USA) coupled to an Orbitrap Fusion™ Tribrid™ Mass Spectrometer (Thermo Scientific, Bremen, Germany) equipped with a nano-electrospray ion source. Peptide mixtures were pre-concentrated onto an Acclaim PepMap 100 C18 trap column (100 μm × 2 cm, Thermo Scientific) and subsequently separated on an EASY-Spray analytical column (25 cm × 75 μm ID) packed with Thermo Scientific Acclaim PepMap RSLC C18 (3 μm, 100 Å). The mobile phase consisted of 0.1% formic acid in water (buffer A) and 0.1% formic acid in acetonitrile (20/80, v/v; buffer B). Chromatographic separation was carried out at a flow rate of 0.300 μL/min.

Phosphoproteomic analysis was performed using 150-minute runs. Following sample loading, the gradient started with 4% buffer B for the first 3 minutes, followed by a linear increase from 5% to 28% over 100 minutes. This was followed by a 10-minute increase to 40% and a subsequent 4-minute step to 95%. The column temperature was maintained at 35°C, and each sample was injected in triplicate with an injection volume of 2 μL. MS analysis was conducted in data-dependent mode (DDA), with a scan range of m/z 375–1500 and a resolution of 120,000. A cycle time of 3 seconds was maintained between master scans. HCD fragmentation was performed with a collision energy of 30 eV, and MS/MS scans were acquired at a resolution of 30,000. The instrument operated in positive polarity mode.

### MS analysis and data processing

All samples underwent label-free quantification (LFQ) analysis using Proteome Discoverer 2.5. The search database was set to Homo sapiens (sp_incl_isoforms, TaxID=9606_and_subtaxonomies, v2023-05-03), with trypsin as the digestion enzyme. The following filters were applied: methionine (M) oxidation, acetylation (protein N-terminus), M-loss, M-loss + acetylation, and phosphorylation (STY) were selected as variable modifications, while carbamidomethyl (C) was designated as a fixed modification. For protein-level analysis, a minimum of two peptides per protein was required. Peptide-level criteria included Xcorr ≥ 2.2, Rank = 1, and high confidence, while at the peptide-spectrum match (PSM) level, Xcorr ≥ 2.2 was applied. Quantification was performed using both unique peptides and razor peptides shared among protein groups. Protein abundance was determined by summing all unique normalized peptide ion abundances across runs.

Data analysis was conducted using Perseus (version 1.6.2.3). Initial filtering steps included the removal of contaminants, reverse entries, and proteins identified solely by a modified peptide. LFQ intensity values were log2-transformed, grouped, and filtered to ensure at least three valid values in at least one condition. Missing values were not imputed (imputation mode set to “none”). For differential expression analysis, protein abundance levels were compared using ANOVA and Student’s T-test, with a significance threshold of 0.05. Proteins exhibiting at least one-fold change were classified as differentially regulated. Gene Ontology (GO) enrichment analysis was performed on dysregulated proteins, focusing on cellular components and molecular functions. A false discovery rate (FDR) threshold of < 0.1 was applied to determine statistical significance. For data visualization, different software packages were used (Microsoft Excel, RStudio, SRplot).

### Plate reader-based secretion assay

HeLa-GH cells stably expressing hGH-FM2-GFP were either mock-treated or transfected with the indicated SMARTpool siRNAs for 48 hours. Following transfection, cells were trypsinized and seeded at a density of 3×10^5^ cells per well into 6-well plates coated with substrates of varying stiffness: 0.5 kPa, 8 kPa, or >GPa. The cells were cultured in complete RPMI-1640 medium. After 24 hours, the medium was replaced with 1 ml of RPMI-1640 medium lacking phenol red, fetal calf serum (FCS), and antibiotics. To induce trafficking of hGH-FM2-GFP, cells were treated with cycloheximide (CHX) at a concentration of 50 μg/ml and D/D solubilizer. Cells were incubated under these conditions for 2 hours. At time 0 and after 2 hours, the culture medium from each well was collected and transferred into a new 6-well plate. GFP fluorescence in the medium was measured using a BioTek Cytation 3 plate reader (Agilent). Subsequently, the remaining cells in each well were washed with 1X PBS, and intracellular GFP fluorescence was measured using the same plate reader. To normalize fluorescence readings for variations in cell number across different conditions, cells were stained with Hoechst 33342. The fluorescence intensity of Hoechst 33342 was used to quantify nuclear fluorescence, which served as a basis for normalizing both intracellular GFP fluorescence and GFP fluorescence detected in the culture medium.

### Immunofluorescence, staining and image analysis

Cells grown on coverslips with elastic modules, 0.5 kPa, 8 kPa or >GPa were fixed with 4% paraformaldehyde and permeabilised with 0.2% saponin, as previously described ^[108]^. Samples were then incubated with selected antibodies (see Table S1) against the antigen of interest at 4°C overnight followed by second antibodies labelled with Alexa Fluor dyes (Invitrogen). The coverslips were then mounted and analysed under a confocal microscope (LSM980; Carl Zeiss; 40× or 63 × oil-immersion objective [1.4 NA]). Images were processed using ImageJ software and the Gaussian blur function, in the case of the inset, was used for presentation where appropriate.function. Excel and GraphPad Prism version 5.0 software were used for data analyses and graphing. Adobe Photoshop was used to adjust the contrast of the images (for presentation only), whereas Adobe Illustrator (Adobe Systems) was used to illustrate figures and draw models.

### Live cell imaging and quantification

For live cell imaging, HeLa-GH were grown on indicated substrate rigidities and then mounted on a Zeiss LSM700 or LSM980 laser scanning microscope under controlled temperature and CO_2_. The images were acquired at regular interval (488 for excitation; PMT: 510–550 nm; 512 × 512 pixels; frame average, 4). Quantitative analysis of hGH-FM2-GFP, TIMP1- and TIMP2-eGFP RUSH trafficking was performed by measuring GFP fluorescence intensity within defined cellular regions. The Golgi region of interest (ROI) was identified based on the characteristic perinuclear accumulation of GFP signal observed at later time points following ligand-induced release from the ER, as previously described ^[16]^. This distinct accumulation pattern served as a spatial reference to retrospectively define the Golgi ROI at earlier time points, ensuring consistent anatomical accuracy throughout the time course.

At each time point, the integrated GFP fluorescence within the Golgi ROI was measured. The extra-Golgi signal was calculated by subtracting the Golgi ROI intensity from the total cellular fluorescence. The ratio of hGH-FM2-GFP integrated intensity in the Golgi versus the extra-Golgi region was then determined and used for data visualization and statistical analysis.

Quantitative analysis of PC-I trafficking was performed by measuring, at each time point, the integrated PC-I fluorescence intensity of the total cell and the TGN fluorescence intensity. The ratio of PC-I integrated intensity to TGN fluorescence intensity was then determined and used for data visualization and statistical analysis.

ImageJ software and Adobe Photoshop were used to adjust the contrast of the movie for presentation only.

### Statistics

Error bars correspond to either standard deviation (SD) or standard error of the mean (SEM) according to the different experiments and as indicated in the figure legends. Statistical evaluations were made by One -way ANOVA, Two-Way ANOVA or Student’s t-test. In all cases, significance was defined as ∗p < 0.05, ∗∗p < 0.01, and ∗∗∗p < 0.001, ∗∗∗∗p < 0.0001, not significant when the significance is not indicated. Statistical analysis was carried out using GraphPad Prism Software.

## Data availability

The mass spectrometry proteomics data have been deposited to the ProteomeXchange Consortium via the PRIDE ^[109]^ partner repository with the dataset identifier PXD061714.

Reviewers can access the dataset by logging in to the PRIDE website using the following account details:

Username

Password: b59DpFvaXrdN

## Author contributions

GS was involved in the planning and execution of most of the experiments, analysis and contributed to the interpretation of data; SF contributed to the execution of experiments and assisted RR in conducting the experiments at the initial phase of the project; ES was involved in the planning and execution of the metabolic labelling experiments; AR contributed to the execution of the experiments; GDB and SC assisted RR in conducting the experiments at the final phase of the project; EJL was involved in cloning and construct preparation; AG and AC contributed to conception and design and interpretation of data; MP was involved in EM sample preparation and acquisition of images; LLDM was involved in the planning of experiments at the initial phase of the project and contributed to the review and editing of the manuscript; RV conducted the experiments with BJ-5ta cells and contributed to the interpretation of data; GDA and SP provided important inputs on the interpretation of the data and contributed to the review and editing of the manuscript; TV contributed to the interpretation of data and editing of the manuscript; GG contributed to the review and editing of the manuscript; ESS provided GBF1 constructs and contributed to conception and design, and interpretation of data and review and editing of the manuscript; LS and PG performed phosphoproteomics MS data analysis and contributed to the writing of the manuscript; LR helped in performing phosphoproteomics MS experiments and contributed to the review and editing of the manuscript; DR contributed to development of the idea, designed and supervised the entire project, and wrote the manuscript; RR developed the idea, designed and supervised the entire project, and wrote the manuscript.

## Acknowledgements

We thank Antonella De Matteis and Advait Subramanian for valuable discussions. Eliana D’Amone for technical assistance. Aurora Rizzo for support in project management. The EUroBioImaging (EUBI) Facility and the PNRR project “SEE LIFE-StrEngthEning the ItaLIan InFrastructure of Euro-bioimaging” at IEOMI-CNR (Naples) for support in EM imaging. The work of RR was supported by the Italian Ministry for Universities and Research (MUR) PRIN-2020 (Prot. 20205B2HZE); by EU funding within the NextGenerationEU – MUR PNRR Extended Partnership initiative on Emerging Infectious Diseases (project no. PE00000007, PE13 INFACT, Spoke 1); by the MUR PRIN-2022 Prot. 2022KEFNPM; and PRIN 2022 PNRR (Prot. P2022CC2N8); by the “Tecnopolo per la medicina di precisione” (TecnoMed Puglia) – Regione Puglia: DGR n.2117 of 21/11/2018. DR acknowledge financial support from MUR PRIN-2022 PNRR (Prot. P2022MMPXH). DR and TV acknowledge the EU funding within the NextGenerationEU – MUR PNRR “National Center for Gene Therapy and Drug based on RNA Technology” (CN00000041). The work was also supported by the Italian Ministry of Research (MUR) in the framework of the National Recovery and Resilience Plan (NRRP), Project Title: “Nano Foundries and Fine Analysis – Digital Infrastructure (NFFA-DI)” (CUP B53C22004310006), funded by the European Union – NextGenerationEU. LLDM acknowledges financial support from ERC-2017-STG (grant number 759959), AIRC-MFAG-2019 (grant number 22902). LR acknowledges financial support from ERC-2019-STG (grant number 850936), the Fondazione Cariplo (grant number 2019-4278), and MUR (PRIN-2022 (Prot. 20205B2HZE), MUR PRIN-2022 Prot. 2022PL44LM; and PRIN 2022 PNRR (Prot. P2022CC2N8_002). R.V. acknowledges the support from the Italian Association for Cancer Research (grant MFAG 2020_25174) and the Italian Ministry of University and Research (MUR, PRIN, 20222MAWZP). ES acknowledges the Fondazione Cariplo (grant number 2022-0438) for the financial support. Models in Figure 1F, Figure 2A, Figure 7, Figure S1E and Figure S3A were created using BioRender.com.

## References

[1] M. Uhlén, M. J. Karlsson, A. Hober, A. S. Svensson, J. Scheffel, D. Kotol, W. Zhong, A. Tebani, L. Strandberg, F. Edfors, E. Sjöstedt, J. Mulder, A. Mardinoglu, A. Berling, S. Ekblad, M. Dannemeyer, S. Kanje, J. Rockberg, M. Lundqvist, M. Malm, A. L. Volk, P. Nilsson, A. Månberg, T. Dodig-Crnkovic, E. Pin, M. Zwahlen, P. Oksvold, K. von Feilitzen, R. S. Häussler, M. G. Hong, C. Lindskog, F. Ponten, B. Katona, J. Vuu, E. Lindström, J. Nielsen, J. Robinson, B. Ayoglu, D. Mahdessian, D. Sullivan, P. Thul, F. Danielsson, C. Stadler, E. Lundberg, G. Bergström, A. Gummesson, B. G. Voldborg, H. Tegel, S. Hober, B. Forsström, J. M. Schwenk, L. Fagerberg, Å. Sivertsson, Sci Signal 2019, 12.

[2] M. Uhlen, M. Uhlén, L. Fagerberg, B. M. Hallström, B. M. Hallstrom, C. Lindskog, P. Oksvold, A. Mardinoglu, A. Sivertsson, A. Sivertsson, C. Kampf, E. Sjöstedt, E. Sjostedt, A. Asplund, I. Olsson, K. Edlund, E. Lundberg, S. Navani, C. A.-K. Szigyarto, J. Odeberg, D. Djureinovic, J. O. Takanen, S. Hober, T. Alm, P. H. Edqvist, P.-H. Edqvist, H. Berling, H. Tegel, J. Mulder, J. Rockberg, P. Nilsson, J. M. Schwenk, M. Hamsten, K. von Feilitzen, K. von Feilitzen, M. Forsberg, L. Persson, F. Johansson, M. Zwahlen, G. von Heijne, G. von Heijne, J. Nielsen, F. Ponten, F. Ponten, Science (1979) 2015, 347.

[3] R. Venditti, C. Wilson, M. A. De Matteis, Exiting the ER: What we know and what we don’t, Vol. 24, 2014.

[4] C. Xu, D. T. W. Ng, Glycosylation-directed quality control of protein folding, Vol. 16, Nature Publishing Group 2015.

[5] Z. Z. Lieu, M. C. Derby, P. A. Gleeson, in The Golgi Apparatus: State of the Art 110 Years after Camillo Golgi’s Discovery, Springer-Verlag Wien 2008, pp. 358–374.

[6] P. Pothukuchi, I. Agliarulo, D. Russo, R. Rizzo, F. Russo, S. Parashuraman, Translation of genome to glycome: role of the Golgi apparatus, Vol. 593, Wiley Blackwell 2019.

[7] H. W. Beams, R. G. Kessel, Int Rev Cytol 1968, 23.

[8] B. S. Glick, A. Nakano, Membrane traffic within the Golgi apparatus, Vol. 25, 2009.

[9] M. A. De Matteis, A. Luini, Exiting the Golgi complex, Vol. 9, 2008.

[10] M. A. De Matteis, A. Luini, New England Journal of Medicine 2011, 365.

[11] A. Gilchrist, C. E. Au, J. Hiding, A. W. Bell, J. Fernandez-Rodriguez, S. Lesimple, H. Nagaya, L. Roy, S. J. C. Gosline, M. Hallett, J. Paiement, R. E. E. Kearney, T. Nilsson, J. J. M. Bergeron, Cell 2006, 127.

[12] H. Farhan, C. Rabouille, Signalling to and from the secretory pathway, Vol. 124, 2011.

[13] A. Luini, M. A. De Matteis, Trends Cell Biol 1993, 3.

[14] M. Bensellam, E. L. Maxwell, J. Y. Chan, J. Luzuriaga, P. K. West, J. C. Jonas, J. E. Gunton, D. R. Laybutt, Diabetologia 2016, 59.

[15] J. E. Campbell, C. B. Newgard, Mechanisms controlling pancreatic islet cell function in insulin secretion, Vol. 22, 2021.

[16] J. Cancino, A. Capalbo, A. DiCampli, M. Giannotta, R. Rizzo, J. E. Jung, R. DiMartino, M. Persico, P. Heinklein, M. Sallese, A. Luini, Dev Cell 2014, 30.

[17] S. Del Giudice, V. De Luca, S. Parizadeh, D. Russo, A. Luini, R. Di Martino, Endogenous and Exogenous Regulatory Signaling in the Secretory Pathway: Role of Golgi Signaling Molecules in Cancer, Vol. 10, 2022.

[18] A. Subramanian, A. Capalbo, N. R. Iyengar, R. Rizzo, A. di Campli, R. Di Martino, M. Lo Monte, A. R. Beccari, A. Yerudkar, C. del Vecchio, L. Glielmo, G. Turacchio, M. Pirozzi, S. G. Kim, P. Henklein, J. Cancino, S. Parashuraman, D. Diviani, F. Fanelli, M. Sallese, A. Luini, Cell 2019, 176.

[19] C. J. Miller, L. A. Davidson, The interplay between cell signalling and mechanics in developmental processes, Vol. 14, 2013.

[20] C. M. Nelson, J. P. Gleghorn, Sculpting organs: Mechanical regulation of tissue development, Vol. 14, 2012.

[21] T. Mammoto, A. Mammoto, D. E. Ingber, Annu Rev Cell Dev Biol 2013, 29.

[22] K. H. Vining, D. J. Mooney, Mechanical forces direct stem cell behaviour in development and regeneration, Vol. 18, 2017.

[23] D. E. Discher, P. Janmey, Y. L. Wang, Tissue cells feel and respond to the stiffness of their substrate, Vol. 310, 2005.

[24] S. Phuyal, F. Baschieri, Endomembranes: Unsung Heroes of Mechanobiology?, Vol. 8, 2020.

[25] S. Phuyal, E. Djaerff, A. Le Roux, M. J. Baker, D. Fankhauser, S. J. Mahdizadeh, V. Reiterer, A. Parizadeh, E. Felder, J. C. Kahlhofer, D. Teis, M. G. Kazanietz, S. Geley, L. Eriksson, P. Roca-Cusachs, H. Farhan, EMBO J 2022, 41.

[26] K. R. Levental, H. Yu, L. Kass, J. N. Lakins, M. Egeblad, J. T. Erler, S. F. T. Fong, K. Csiszar, A. Giaccia, W. Weninger, M. Yamauchi, D. L. Gasser, V. M. Weaver, Cell 2009, 139.

[27] J. K. Mouw, Y. Yui, L. Damiano, R. O. Bainer, J. N. Lakins, I. Acerbi, G. Ou, A. C. Wijekoon, K. R. Levental, P. M. Gilbert, E. S. Hwang, Y. Y. Chen, V. M. Weaver, Nat Med 2014, 20.

[28] J. C. Friedland, M. H. Lee, D. Boettiger, Science (1979) 2009, 323.

[29] D. Lachowski, C. Matellan, S. Gopal, E. Cortes, B. K. Robinson, A. Saiani, A. F. Miller, M. M. Stevens, A. E. Del Río Hernández, ACS Nano 2022, 16.

[30] R. Masuzaki, R. Tateishi, H. Yoshida, T. Sato, T. Ohki, T. Goto, H. Yoshida, S. Sato, Y. Sugioka, H. Ikeda, S. Shiina, T. Kawabe, M. Omata, Hepatol Int 2007, 1.

[31] F. Liu, J. D. Mih, B. S. Shea, A. T. Kho, A. S. Sharif, A. M. Tager, D. J. Tschumperlin, Journal of Cell Biology 2010, 190.

[32] S. Mueller, Hepat Med 2010.

[33] C. Loebel, R. L. Mauck, J. A. Burdick, Nat Mater 2019, 18.

[34] C. Loebel, M. Y. Kwon, C. Wang, L. Han, R. L. Mauck, J. A. Burdick, Adv Funct Mater 2020, 30.

[35] A. M. Tartakoff, EMBO J 1986, 5.

[36] E. J. Arenas, A. Martínez-Sabadell, I. Rius Ruiz, M. Román Alonso, M. Escorihuela, A. Luque, C. A. Fajardo, A. Gros, C. Klein, J. Arribas, Nat Commun 2021, 12.

[37] D. Lachowski, E. Cortes, A. Rice, D. Pinato, K. Rombouts, A. del Rio Hernandez, Sci Rep 2019, 9.

[38] D. Pankova, Y. Chen, M. Terajima, M. J. Schliekelman, B. N. Baird, M. Fahrenholtz, L. Sun, B. J. Gill, T. J. Vadakkan, M. P. Kim, Y. H. Ahn, J. D. Roybal, X. Liu, E. R. P. Cuentas, J. Rodriguez, I. I. Wistuba, C. J. Creighton, D. L. Gibbons, J. M. Hicks, M. E. Dickinson, J. L. West, K. J. Grande-Allen, S. M. Hanash, M. Yamauchi, J. M. Kurie, Molecular Cancer Research 2016, 14.

[39] A. A. Mironov, G. V. Beznoussenko, P. Nicoziani, O. Martella, A. Trucco, H. S. Kweon, D. Di Giandomenico, R. S. Polishchuk, A. Fusella, P. Lupetti, E. G. Berger, W. J. C. Geerts, A. J. Koster, K. N. J. Burger, A. Luini, Journal of Cell Biology 2001, 155.

[40] D. E. Gordon, L. M. Bond, D. A. Sahlender, A. A. Peden, Traffic 2010, 11.

[41] J. Bass, C. Turck, M. Rouard, D. F. Steiner, Proc Natl Acad Sci U S A 2000, 97.

[42] G. Boncompain, S. Divoux, N. Gareil, H. De Forges, A. Lescure, L. Latreche, V. Mercanti, F. Jollivet, G. Raposo, F. Perez, Nat Methods 2012, 9.

[43] M. J. Paszek, N. Zahir, K. R. Johnson, J. N. Lakins, G. I. Rozenberg, A. Gefen, C. A. Reinhart-King, S. S. Margulies, M. Dembo, D. Boettiger, D. A. Hammer, V. M. Weaver, Cancer Cell 2005, 8.

[44] K. R. Levental, H. Yu, L. Kass, J. N. Lakins, M. Egeblad, J. T. Erler, S. F. T. Fong, K. Csiszar, A. Giaccia, W. Weninger, M. Yamauchi, D. L. Gasser, V. M. Weaver, Cell 2009, 139, 891.

[45] P. Romani, I. Brian, G. Santinon, A. Pocaterra, M. Audano, S. Pedretti, S. Mathieu, M. Forcato, S. Bicciato, J. B. Manneville, N. Mitro, S. Dupont, Nat Cell Biol 2019, 21.

[46] K. S. Matlin, K. Simons, Cell 1983, 34.

[47] D. J. Klionsky, H. Abeliovich, P. Agostinis, D. K. Agrawal, G. Aliev, D. S. Askew, M. Baba, E. H. Baehrecke, B. A. Bahr, A. Ballabio, B. A. Bamber, D. C. Bassham, E. Bergamini, X. Bi, M. Biard-Piechaczyk, J. S. Blum, D. E. Bredesen, J. L. Brodsky, J. H. Brumell, U. T. Brunk, W. Bursch, N. Camougrand, E. Cebollero, F. Cecconi, Y. Chen, L. S. Chin, A. Choi, C. T. Chu, J. Chung, P. G. H. Clarke, R. S. B. Clark, S. G. Clarke, C. Clavé, J. L. Cleveland, P. Codogno, M. I. Colombo, A. Cotomontes, J. M. Cregg, A. M. Cuervo, J. Debnath, F. Demarchi, P. B. Dennis, P. A. Dennis, V. Deretic, R. J. Devenish, F. Di Sano, J. F. Dice, M. DiFiglia, S. Dinesh-Kumar, C. W. Distelhorst, M. Djavaheri-Mergny, F. C. Dorsey, W. Dröge, M. Dron, W. A. Dunn, M. Duszenko, N. T. Eissa, Z. Elazar, A. Esclatine, E. L. Eskelinen, L. Fésüs, K. D. Finley, J. M. Fuentes, J. Fueyo, K. Fujisaki, B. Galliot, F. B. Gao, D. A. Gewirtz, S. B. Gibson, A. Gohla, A. L. Goldberg, R. Gonzalez, C. González-Estévez, S. Gorski, R. A. Gottlieb, D. Häussinger, Y. W. He, K. Heidenreich, J. A. Hill, M. Høyer-Hansen, X. Hu, W. P. Huang, A. Iwasaki, M. Jäättelä, W. T. Jackson, X. Jiang, S. Jin, T. Johansen, J. U. Jung, M. Kadowaki, C. Kang, A. Kelekar, D. H. Kessel, J. A. K. W. Kiel, P. K. Hong, A. Kimchi, T. J. Kinsella, K. Kiselyov, K. Kitamoto, E. Knecht, M. Komatsu, E. Kominami, S. Kondo, A. L. Kovács, G. Kroemer, C. Y. Kuan, R. Kumar, M. Kundu, J. Landry, M. Laporte, W. Le, H. Y. Lei, M. J. Lenardo, B. Levine, A. Lieberman, K. L. Lim, F. C. Lin, W. Liou, L. F. Liu, G. Lopez-Berestein, C. López-Otín, B. Lu, K. F. Macleod, W. Malorni, W. Martinet, K. Matsuoka, J. Mautner, A. J. Meijer, A. Meléndez, P. Michels, G. Miotto, W. P. Mistiaen, N. Mizushima, B. Mograbi, I. Monastyrska, M. N. Moore, P. I. Moreira, Y. Moriyasu, T. Motyl, C. Münz, L. O. Murphy, N. I. Naqvi, T. P. Neufeld, I. Nishino, R. A. Nixon, T. Noda, B. Nürnberg, M. Ogawa, N. L. Oleinick, L. J. Olsen, B. Ozpolat, S. Paglin, G. E. Palmer, I. Papassideri, M. Parkes, D. H. Perlmutter, G. Perry, M. Piacentini, R. Pinkas-Kramarski, M. Prescott, T. Proikascezanne, N. Raben, A. Rami, F. Reggiori, B. Rohrer, D. C. Rubinsztein, K. M. Ryan, J. Sadoshima, H. Sakagami, Y. Sakai, M. Sandri, C. Sasakawa, M. Sass, C. Schneider, P. O. Seglen, O. Seleverstov, J. Settleman, J. J. Shacka, I. M. Shapiro, A. Sibirny, E. C. M. Silva-Zacarin, H. U. Simon, C. Simone, A. Simonsen, M. A. Smith, K. Spanel-Borowski, V. Srinivas, M. Steeves, H. Stenmark, P. E. Stromhaug, C. S. Subauste, S. Sugimoto, D. Sulzer, T. Suzuki, M. S. Swanson, I. Tabas, F. Takeshita, N. J. Talbot, Z. Tallóczy, K. Tanaka, K. Tanaka, I. Tanida, G. S. Taylor, J. P. Taylor, A. Terman, G. Tettamanti, C. B. Thompson, M. Thumm, A. M. Tolkovsky, S. A. Tooze, R. Truant, L. V. Tumanovska, Y. Uchiyama, T. Ueno, N. L. Uzcátegui, I. Van Der Klei, E. C. Vaquero, T. Vellai, M. W. Vogel, H. G. Wang, P. Webster, J. W. Wiley, Z. Xi, G. Xiao, J. Yahalom, J. M. Yang, G. Yap, X. M. Yin, T. Yoshimori, L. Yu, Z. Yue, M. Yuzaki, O. Zabirnyk, X. Zheng, X. Zhu, R. L. Deter, Guidelines for the use and interpretation of assays for monitoring autophagy in higher eukaryotes, Vol. 4, 2008.

[48] H. Palokangas, M. Ying, K. Väänänen, J. Saraste, Mol Biol Cell 1998, 9.

[49] K. Burridge, Focal adhesions: a personal perspective on a half century of progress, Vol. 284, 2017.

[50] P. Kanchanawong, D. A. Calderwood, Organization, dynamics and mechanoregulation of integrin-mediated cell–ECM adhesions, Vol. 24, 2023.

[51] F. G. Giancotti, G. Tarone, in Annual Review of Cell and Developmental Biology, Vol. 19, 2003.

[52] M. A. Schwartz, M. D. Schaller, M. H. Ginsberg, Integrins: Emerging paradigms of signal transduction, Vol. 11, 1995.

[53] L. Koudelková, J. Brábek, D. Rosel, Src kinase: Key effector in mechanosignalling, Vol. 131, 2021.

[54] S. K. Mitra, D. D. Schlaepfer, Integrin-regulated FAK-Src signaling in normal and cancer cells, Vol. 18, 2006.

[55] R. K. Assoian, E. A. Klein, Growth control by intracellular tension and extracellular stiffness, Vol. 18, 2008.

[56] F. Haun, S. Neumann, L. Peintner, K. Wieland, J. Habicht, C. Schwan, K. Østevold, M. M. Koczorowska, M. Biniossek, M. Kist, H. Busch, M. Boerries, R. J. Davis, U. Maurer, O. Schilling, K. Aktories, C. Borner, Nat Commun 2018, 9.

[57] J. Sun, X. Wang, B. Tang, H. Liu, M. Zhang, Y. Wang, F. Ping, J. Ding, A. Shen, M. Geng, Theranostics 2018, 8.

[58] F. Bard, L. Mazelin, C. Péchoux-Longin, V. Malhotra, P. Jurdic, Journal of Biological Chemistry 2003, 278.

[59] M. Giannotta, C. Ruggiero, M. Grossi, J. Cancino, M. Capitani, T. Pulvirenti, G. M. L. Consoli, C. Geraci, F. Fanelli, A. Luini, M. Sallese, EMBO Journal 2012, 31.

[60] M. Fu, Y. Hu, T. Lan, K.-L. Guan, T. Luo, M. Luo, Signal Transduct Target Ther 2024, 9.

[61] T. Szul, R. Grabski, S. Lyons, Y. Morohashi, S. Shestopal, M. Lowe, E. Sztul, J Cell Sci 2007, 120.

[62] R. K. Assoian, E. A. Klein, Growth control by intracellular tension and extracellular stiffness, Vol. 18, 2008.

[63] M. Giannotta, C. Ruggiero, M. Grossi, J. Cancino, M. Capitani, T. Pulvirenti, G. M. L. Consoli, C. Geraci, F. Fanelli, A. Luini, M. Sallese, EMBO Journal 2012, 31, 2869.

[64] G. A. Belov, Q. Feng, K. Nikovics, C. L. Jackson, E. Ehrenfeld, PLoS Pathog 2008, 4.

[65] K. Walton, T. J. Nawara, A. R. Angermeier, H. Rosengrant, E. Lee, B. Wynn, E. Victorova, G. Belov, E. Sztul, Sci Rep 2023, 13.

[66] C. Ford, C. G. Burd, Mol Biol Cell 2022, 33.

[67] T. Pelaseyed, G. C. Hansson, J Cell Sci 2011, 124.

[68] M. Lopes-da-Silva, J. J. McCormack, J. J. Burden, K. J. Harrison-Lavoie, F. Ferraro, D. F. Cutler, Dev Cell 2019, 49.

[69] L. Mao, N. Li, Y. Guo, X. Xu, L. Gao, Y. Xu, L. Zhou, W. Liu, J Cell Sci 2013, 126.

[70] T. Miyamoto, N. Oshiro, K. I. Yoshino, A. Nakashima, S. Eguchi, M. Takahashi, Y. Ono, U. Kikkawa, K. Yonezawa, Journal of Biological Chemistry 2008, 283.

[71] X. Xu, Y. Fang, S. Nowsheen, Y. X. Li, Z. Lou, M. Deng, Genes Dis 2024, 11.

[72] A. C. Ferretti, F. Hidalgo, F. M. Tonucci, E. Almada, A. Pariani, M. C. Larocca, C. Favre, Sci Rep 2019, 9.

[73] J. Winkler, A. Abisoye-Ogunniyan, K. J. Metcalf, Z. Werb, Concepts of extracellular matrix remodelling in tumour progression and metastasis, Vol. 11, 2020.

[74] J. Jung, M. M. Khan, J. Landry, A. Halavatyi, P. Machado, M. Reiss, R. Pepperkok, Journal of Cell Biology 2022, 221.

[75] S. Phuyal, E. Djaerff, A. Le Roux, M. J. Baker, D. Fankhauser, S. J. Mahdizadeh, V. Reiterer, A. Parizadeh, E. Felder, J. C. Kahlhofer, D. Teis, M. G. Kazanietz, S. Geley, L. Eriksson, P. Roca-Cusachs, H. Farhan, EMBO J 2022, 41.

[76] J. Lowery, T. Szul, M. Styers, Z. Holloway, V. Oorschot, J. Klumperman, E. Sztul, Journal of Biological Chemistry 2013, 288.

[77] A. Stockhammer, P. Adarska, V. Natalia, A. Heuhsen, A. Klemt, G. Bregu, S. Harel, C. Rodilla-Ramirez, C. Spalt, E. Özsoy, P. Leupold, A. Grindel, E. Fox, J. O. Mejedo, A. Zehtabian, H. Ewers, D. Puchkov, V. Haucke, F. Bottanelli, Nat Cell Biol 2024.

[78] J. T. Parsons, A. R. Horwitz, M. A. Schwartz, Cell adhesion: Integrating cytoskeletal dynamics and cellular tension, Vol. 11, 2010.

[79] V. Vogel, M. P. Sheetz, Cell fate regulation by coupling mechanical cycles to biochemical signaling pathways, Vol. 21, 2009.

[80] K. R. Levental, H. Yu, L. Kass, J. N. Lakins, M. Egeblad, J. T. Erler, S. F. T. Fong, K. Csiszar, A. Giaccia, W. Weninger, M. Yamauchi, D. L. Gasser, V. M. Weaver, Cell 2009, 139, 891.

[81] M. M. Mihaylova, R. J. Shaw, The AMPK signalling pathway coordinates cell growth, autophagy and metabolism, Vol. 13, 2011.

[82] M. Zhao, D. Finlay, E. Kwong, R. Liddington, B. Viollet, N. Sasaoka, K. Vuori, Cell Signal 2022, 89.

[83] A. Stockhammer, P. Adarska, V. Natalia, A. Heuhsen, A. Klemt, G. Bregu, S. Harel, C. Rodilla-Ramirez, C. Spalt, E. Özsoy, P. Leupold, A. Grindel, E. Fox, J. O. Mejedo, A. Zehtabian, H. Ewers, D. Puchkov, V. Haucke, F. Bottanelli, Nat Cell Biol 2024.

[84] C. C. Dibble, L. C. Cantley, Regulation of mTORC1 by PI3K signaling, Vol. 25, 2015.

[85] F. Y. Lee, Y. Y. Zhen, C. M. Yuen, R. Fan, Y. T. Chen, J. J. Sheu, Y. L. Chen, C. J. Wang, C. K. Sun, H. K. Yip, Am J Transl Res 2017, 9.

[86] D. M. Gwinn, D. B. Shackelford, D. F. Egan, M. M. Mihaylova, A. Mery, D. S. Vasquez, B. E. Turk, R. J. Shaw, Mol Cell 2008, 30.

[87] R. J. Shaw, N. Bardeesy, B. D. Manning, L. Lopez, M. Kosmatka, R. A. DePinho, L. C. Cantley, Cancer Cell 2004, 6.

[88] K. Inoki, T. Zhu, K. L. Guan, Cell 2003, 115.

[89] C. Justicia, J. Panés, S. Solé, Á. Cervera, R. Deulofeu, Á. Chamorro, A. M. Planas, Journal of Cerebral Blood Flow and Metabolism 2003, 23.

[90] A. Ivanoff, J. Ivanoff, K. Hultenby, K. G. Sundqvist, Clin Exp Metastasis 1999, 17.

[91] S. Rangarajan, N. B. Bone, A. A. Zmijewska, S. Jiang, D. W. Park, K. Bernard, M. L. Locy, S. Ravi, J. Deshane, R. B. Mannon, E. Abraham, V. Darley-Usmar, V. J. Thannickal, J. W. Zmijewski, Nat Med 2018, 24.

[92] D. Cheng, Q. Xu, Y. Wang, G. Li, W. Sun, D. Ma, S. Zhou, Y. Liu, L. Han, C. Ni, J Transl Med 2021, 19.

[93] Z. Liang, T. Li, S. Jiang, J. Xu, W. di, Z. Yang, W. Hu, Y. Yang, AMPK: A novel target for treating hepatic fibrosis, Vol. 8, 2017.

[94] S. Jiang, T. Li, Z. Yang, W. Yi, S. Di, Y. Sun, D. Wang, Y. Yang, AMPK orchestrates an elaborate cascade protecting tissue from fibrosis and aging, Vol. 38, 2017.

[95] Q. Liu, Y. Zhou, J. D. Cogan, D. B. Mitchell, Q. Sheng, S. Zhao, Y. Bai, K. K. Ciombor, C. M. Sabusap, M. M. Malabanan, C. R. Markin, K. Douglas, G. Ding, N. E. Banovich, D. A. Nickerson, E. E. Blue, M. J. Bamshad, K. K. Brown, D. A. Schwartz, J. A. Phillips, R. Martinez-Barricarte, M. L. Salisbury, Y. Shyr, J. E. Loyd, J. A. Kropski, T. S. Blackwell, Am J Respir Crit Care Med 2023, 207, 1345.

[96] P. A. Janmey, D. A. Fletcher, C. A. Reinhart-King, Physiol Rev 2020, 100.

[97] G. Huang, F. Li, X. Zhao, Y. Ma, Y. Li, M. Lin, G. Jin, T. J. Lu, G. M. Genin, F. Xu, Functional and Biomimetic Materials for Engineering of the Three-Dimensional Cell Microenvironment, Vol. 117, 2017.

[98] L. F. Mattner, Z. Zeng, C. H. Mayr, M. Ansari, X. Wei, S. Asgharpour, A. A. Wasik, N. Kneidinger, M.-G. Stoleriu, J. Behr, J. Polleux, A. Ö. Yildirim, G. Burgstaller, M. Mann, H. B. Schiller, bioRxiv 2023.

[99] P. Lu, K. Takai, V. M. Weaver, Z. Werb, Cold Spring Harb Perspect Biol 2011, 3.

[100] A. A. Gibb, M. P. Lazaropoulos, J. W. Elrod, Myofibroblasts and fibrosis: Mitochondrial and metabolic control of cellular differentiation, Vol. 127, 2020.

[101] C. Bonnans, J. Chou, Z. Werb, Remodelling the extracellular matrix in development and disease, Vol. 15, 2014.

[102] R. García-Mata, T. Szul, C. Alvarez, E. Sztul, Mol Biol Cell 2003, 14.

[103] C. Loebel, A. M. Saleh, K. R. Jacobson, R. Daniels, R. L. Mauck, S. Calve, J. A. Burdick, Metabolic labeling of secreted matrix to investigate cell–material interactions in tissue engineering and mechanobiology, Vol. 17, 2022.

[104] H. P. Hauri, C. Appenzeller, F. Kuhn, O. Nufer, Lectins and traffic in the secretory pathway, Vol. 476, 2000.

[105] J. Füllekrug, P. Scheiffele, K. Simons, J Cell Sci 1999, 112.

[106] A. Mironov, G. V. Beznoussenko, A. Trucco, P. Lupetti, J. D. Smith, W. J. C. Geerts, A. J. Koster, K. N. J. Burger, M. E. Martone, T. J. Deerinck, M. H. Ellisman, A. Luini, Dev Cell 2003, 5.

[107] A. Trucco, R. S. Polischuck, O. Martella, A. Di Pentima, A. Fusella, D. Di Giandomenico, E. San Pietro, G. V. Beznoussenko, E. V. Polischuk, M. Baldassarre, R. Buccione, W. J. C. Geerts, A. J. Koster, K. N. J. Burger, A. A. Mironov, A. Luini, Nat Cell Biol 2004, 6.

[108] R. Rizzo, D. Russo, K. Kurokawa, P. Sahu, B. Lombardi, D. Supino, M. A. Zhukovsky, A. Vocat, P. Pothukuchi, V. Kunnathully, L. Capolupo, G. Boncompain, C. Vitagliano, F. Zito Marino, G. Aquino, D. Montariello, P. Henklein, L. Mandrich, G. Botti, H. Clausen, U. Mandel, T. Yamaji, K. Hanada, A. Budillon, F. Perez, S. Parashuraman, Y. A. Hannun, A. Nakano, D. Corda, G. D’Angelo, A. Luini, EMBO J 2021, 40.

[109] Y. Perez-Riverol, C. Bandla, D. J. Kundu, S. Kamatchinathan, J. Bai, S. Hewapathirana, N. S. John, A. Prakash, M. Walzer, S. Wang, J. A. Vizcaíno, Nucleic Acids Res 2025, 53, D543.

