## Supplementary material for "Mechanical Cues Regulate Cargo Sorting and Export at the Golgi": Revised_Supporting information_PD.pdf

### Supplementary figures

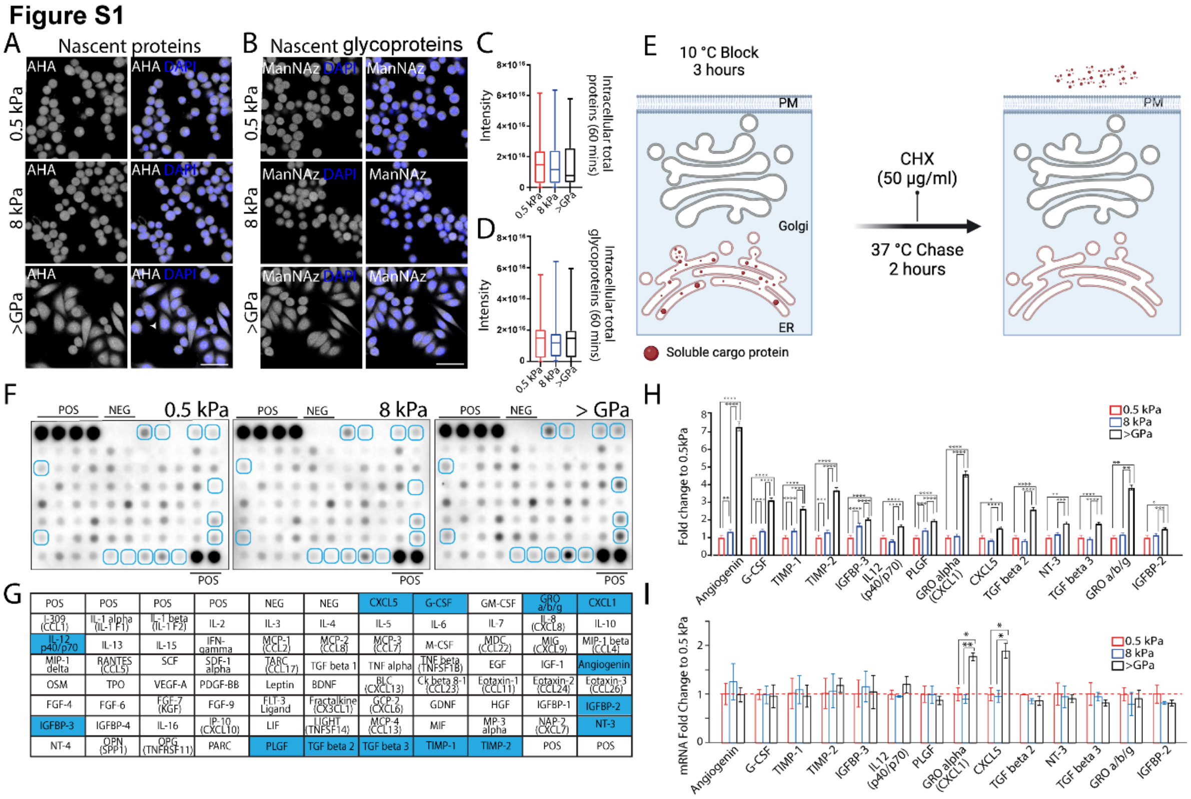


**Figure S1. Transcriptional profiling of stiffness-dependent secreted cargoes.**

**A, B.** HeLa cells were grown on the indicated substrate rigidities coated with 10 µg/ml human fibronectin. The azide-modified methionine analogue (AHA, [A]) or a sugar analogue (ManNAz, [B]) was added to the culture media for 1 hours. Then, cells were fixed end permeabilized prior to proceed with bio-orthogonal strain-promoted cyclo-addition with fluorophore-modified cyclooctyne (DBCO Cy5). Representative images of total intracellular nascent proteins (AHA, [A]) and glycoproteins (ManNAz, [B])are shown. DAPI (blue), AHA or ManNAz (white). Scale bar, 50 µm. **C**, **D**. Quantification of the total intracellular fluorescence intensity of proteins (C) from (A) and glycoproteins (D) from (B) (data are means ± SEM derived from 2 biological replicates, in which cumulatively >2500 cells have been quantified for each condition. **E**. Synchronisation protocol for protein secretion (created with BioRender.com). HeLa cells were incubated for 3 hours at 10°C, then shifted to 37°C for 2 hours in the presence of cycloheximide (CHX, 50 µg/ml). **F**. Synchronised cytokine secretion of cell seeded on silicone substrates with elastic modulus of 0.5 kPa, 8 kPa, or >GPa. HeLa cells grown on indicated substrate rigidities were subjected to protein synchronisation protocol as in (E). After 2 hours of chase, the medium was collected and used for cytokine array analysis according to the manufacturer instructions. The factors showing an increase from soft to stiff (0.5 kPa ≤ 8 kPa ≤ >GPa) are indicated by the blue boxes. POS, positive controls consisting of controlled amount of biotinylated antibody printed on to the array; NEG, negative controls consisting of Buffer printed on to the array (no antibodies). **G**. Schematic representation of the cytokine spot positions on the membrane. Rectangles filled with blue background represents the proteins indicated by the blue boxes in (F). **H.** Densitometric quantification of the blot in (F) (blue boxes in [F and G]) (data are means of 2 experiments ± SD. *p< 0.05, **p<0.01, ***p<0.001, ****p<0.0001 [One-way ANOVA]). **I**. Cytokines mRNA expression on different substrate rigidities. mRNA levels of indicated genes were evaluated in HeLa cells cultured on different substrate rigidities, normalised to control hypoxanthine phosphoribosyltransferase 1 (HPRT1) mRNA and presented relative to 0.5 kPa sample (data are means of at least 3 independent experiments ± SD. *p<0.05, **p<0.01, [One-way ANOVA]).

### Figure S2

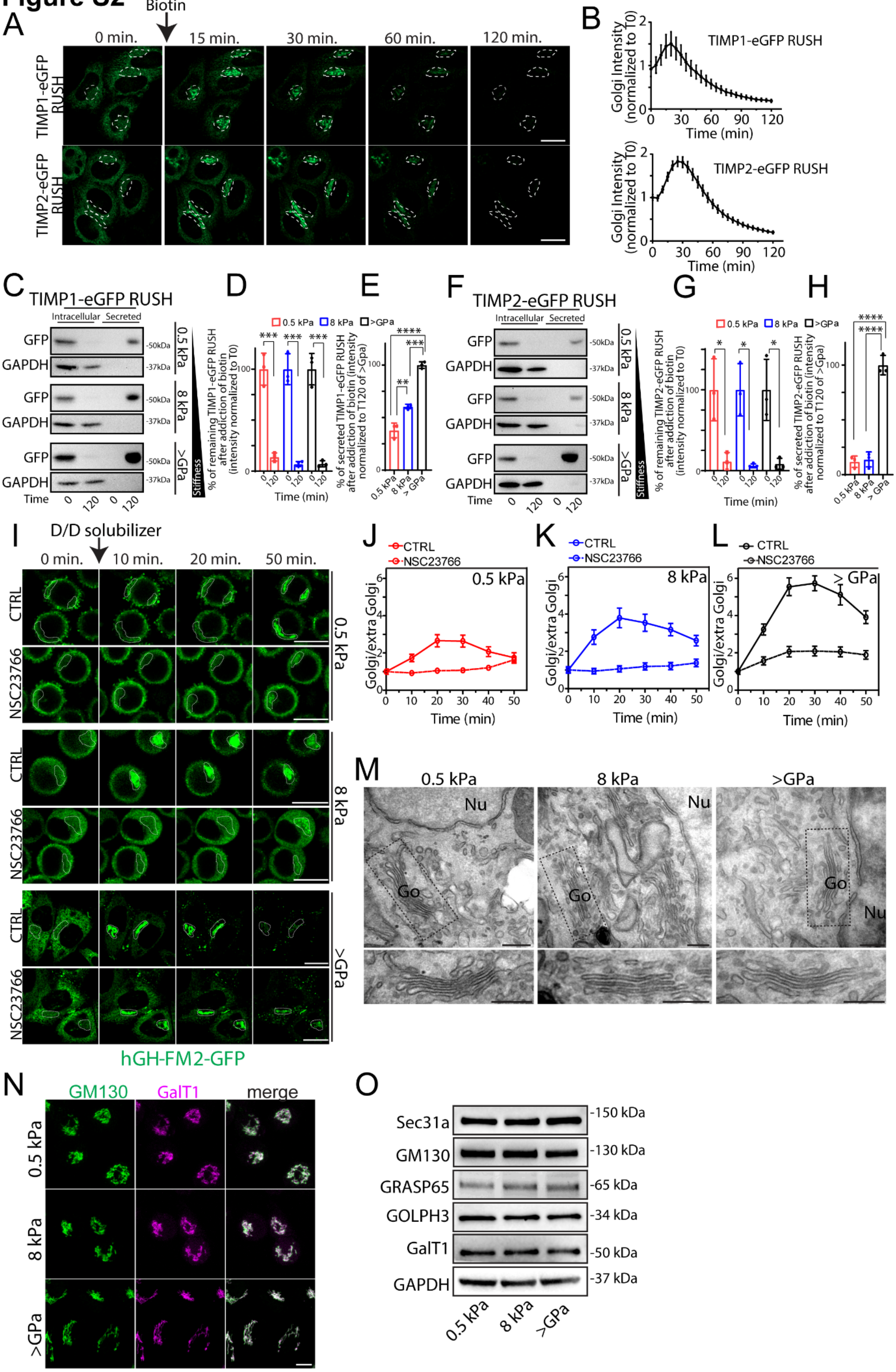
A
Biotin
0 min.
15 min.
30 min.
60 min.
120 min.
TIMP1-eGFP RUSH
TIMP2-eGFP RUSH
B
Golgi Intensity (normalized to T0)
TIMP1-eGFP RUSH
TIMP2-eGFP RUSH
Time (min)
C
TIMP1-eGFP RUSH
Intracellular
Secreted
GFP
GAPDH
-50kDa
-37kDa
0.5 kPa
8 kPa
>GPa
Stiffness
Time
D
0.5 kPa
8 kPa
>GPa
% of remaining TIMP1-eGFP RUSH after addiction of biotin (intensity normalized to T0)
Time (min)
E
% of secreted TIMP1-eGFP RUSH after addiction of biotin (intensity normalized to T120 of >Gpa)
F
TIMP2-eGFP RUSH
G
% of remaining TIMP2-eGFP RUSH after addiction of biotin (intensity normalized to T0)
H
% of secreted TIMP2-eGFP RUSH after addiction of biotin (intensity normalized to T120 of >Gpa)
I
D/D solubilizer
0 min.
10 min.
20 min.
50 min.
CTRL
NSC23766
hGH-FM2-GFP
J
K
L
CTRL
NSC23766
Golgi/extra Golgi
Time (min)
M
0.5 kPa
8 kPa
>GPa
Nu
Go
N
GM130
GalT1
merge
O
Sec31a
GM130
GRASP65
GOLPH3
GalT1
GAPDH
-150 kDa
-130 kDa
-65 kDa
-34 kDa
-50 kDa
-37 kDa
0.5 kPa
8 kPa
>GPa

**Figure S2. Stiffness-driven regulation of ER-to-Golgi protein transport occurs independently of Rac1 and without altering Golgi morphology.**

**A.** Assessment of the synchronized transport of the TIMP1-eGFP-RUSH (upper panel) or TIMP2-eGFP-RUSH (bottom panel) in HeLa cells grown on indicated rigidities. HeLa cells transfected with TIMP1-eGFP-RUSH or TIMP2-eGFP-RUSH were grown on stiff substrate (>GPa). After 24 hours, cells were subjected to biotin, (80 μM) to induce the release of cargoes from the ER, in the presence of cycloheximide (CHX, 50 μg/ml), and the transport of TIMP1-eGFP-RUSH or TIMP2-eGFP-RUSH visualised by video microscopy. Micrographs showing cells at the indicated trafficking time points. Dashed lines indicate the Golgi region. Scale bar, 20 μm. **B.** Quantification of the experiment in (A) shows the TIMP1-eGFP-RUSH (upper panel) or TIMP2-eGFP-RUSH (bottom panel) fluorescence intensity in the Golgi region at different time points after biotin and CHX addition. Data are means ± SEM. **C.** HeLa cells expressing TIMP1-eGFP-RUSH were grown on substrate of varying rigidities (0.5 kPa, 8 kPa and >GPa), and treated as in (A, upper panel). The cell lysate (intracellular) and the medium (secreted) were collected at indicated timepoint and analysed by SDS-PAGE (20% of the total intracellular lysate and 100% of the total secreted was loaded, respectively) followed by Western blot. **D, E.** Densitometric quantification of (C) shows intracellular (D) or secreted (E) levels of TIMP1-eGFP-RUSH after 2 hours (data are mean of at least 3 experiments ± SD. $^{**}p<0.01$, $^{***}p<0.001$, $^{****}p<0.0001$ [Two-way ANOVA]). **F.** HeLa cells expressing TIMP2-eGFP-RUSH were grown on substrate of varying rigidities (0.5 kPa, 8 kPa and >GPa), and treated as in (A, bottom panel). The cell lysate (intracellular) and the medium (secreted) were collected at indicated timepoint and analysed by SDS-PAGE (20% of the total intracellular lysate and 100% of the total secreted was loaded, respectively) followed by Western blot. **G, H.** Densitometric quantification of (F) shows intracellular (G) or secreted (H) levels of TIMP2-eGFP-RUSH after 2 hours (data are mean of at least 3 experiments ± SD.

*$p<0.05$, ****$p<0.0001$ [One-way ANOVA]). **I.** Visualization of synchronized hGH-FM2-GFP transport in HeLa-GH cells treated with the Rac1 inhibitor NSC23766. HeLa-GH cells were cultured on 2D silicone substrates with defined elastic moduli (0.5 kPa, 8 kPa, and >GPa). After 24 hours, cells were pre-treated with DMSO (CTRL) or 50 µM NSC23766 (Rac1 inhibitor) for 4 hours, with the drugs maintained during hGH-FM2-GFP trafficking, induced by the addition of D/D (1 µM) and CHX (50 µg/ml). Cells were imaged at the indicated time points. Micrographs show representative cells at each trafficking time point. Dashed lines indicate the Golgi region. Scale bar, 20 µm. **J-L**. Quantification of the experiment in (I) showing Golgi arrival kinetics of hGH-FM2-GFP (n=11 cells measured for each condition; data are means ± SEM. **M**. HeLa-GH were seeded on indicated substrate rigidities for 24 hours, fixed, processed and then examined by electron microscopy. Delineated regions of interest (boxed areas) are enlarged below. Nucleus (Nu); Golgi (Gu). Scale bar, 200 nm. **N**. Fluorescence microscopy images (maximum intensity projection) of HeLa cells grown on indicated substrate rigidities for 24 hours, fixed, and then processed for immunofluorescence staining: GM130 (green), GalT1(magenta). Scale bar, 10 µm. **O**. Protein lysates from HeLa cells seeded on indicated substrate rigidities for 24 hours, lysed and then processed for SDS-PAGE and Western blot with the indicated antibodies. Data are representative of at least 3 independent experiments.

Figure S3

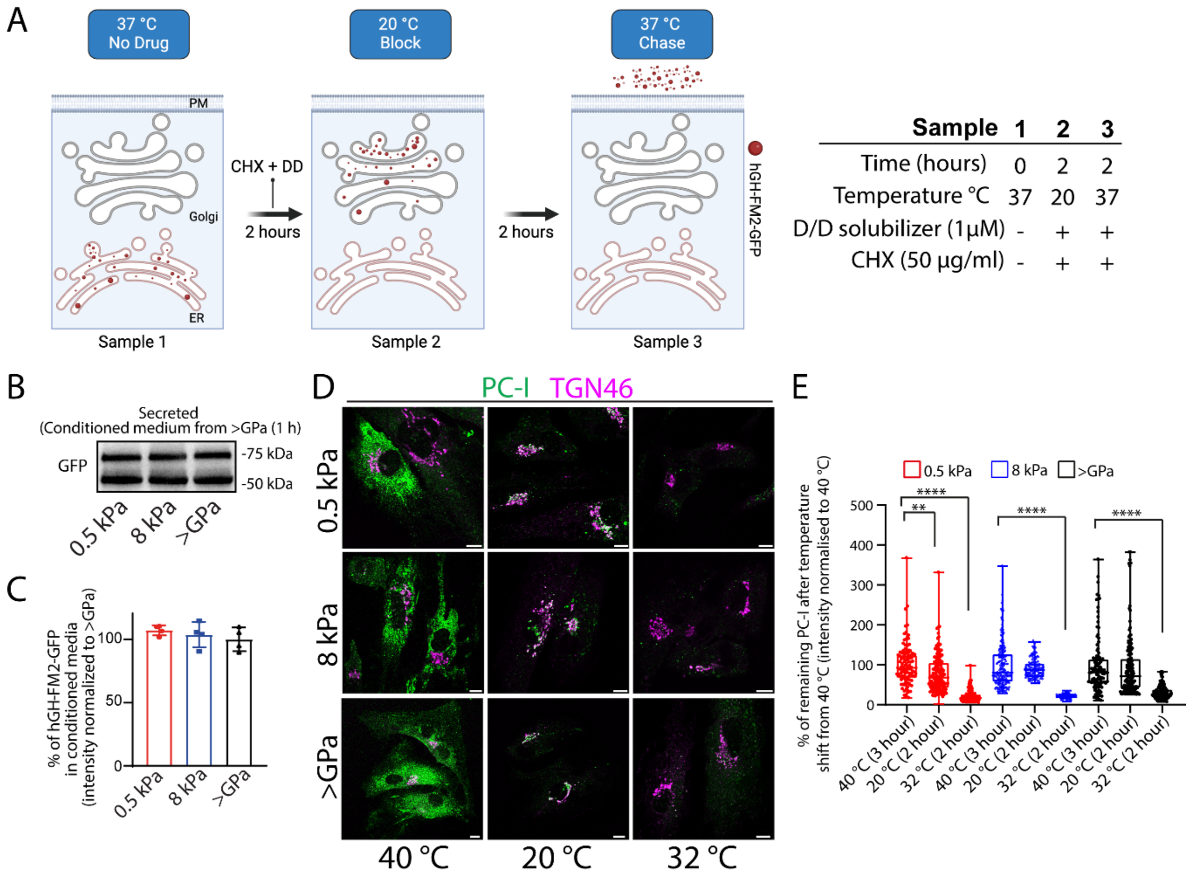


**Figure S3. Stiffness-induced degradation of hGH-FM2-GFP cargo does not occur extracellularly.**

**A**. Schematic representation of 20°C temperature traffic block assay to synchronize ER to Golgi and Golgi to PM transport of hGH-FM2-GFP (created with BioRender.com). In sample 1, hGH is retained in the ER across all stiffness conditions, and no degradation of the cargo is observed. In sample 2, hGH is retained in the Golgi across all stiffnesses, and no degradation is detected. In sample 3, hGH is allowed to exit the Golgi and traffic further for 2 hours with a decrease in its intracellular cargo levels under all stiffness conditions. However, while cells on stiff substrates secrete the cargo (as evidenced by its detection in the medium), cells on soft substrates do not show a corresponding increase in secretion. This suggests that the cargo is being degraded, and that the degradation step occurs post-Golgi. **B.** Conditioned medium containing secreted hGH-FM2-GFP was collected from HeLa-GH cells cultured on stiff substrates (>GPa) after 2 hours of trafficking activation. This medium was then applied to recipient HeLa-GH cells grown on 0.5 kPa, 8 kPa, and >GPa substrates, in which trafficking had also been previously activated for 2 hours. After replacing the medium with the conditioned medium, cells were incubated for an additional 1 hour. The medium (secreted) were collected at indicated timepoint and analysed by SDS-PAGE (100% of the total secreted was loaded, respectively) followed by Western blot. **C.** Densitometric quantification of (B) (data are mean of at least 3 experiments ± SD. **D.** BJ5ta cells were seeded on indicated substrate rigidities and kept for 3-hour incubation at 40°C in the absence of ascorbic acid. Then, cells were shifted to 20°C in the presence of ascorbic acid (100 µg/ml) for 2 hours followed by incubation at 32°C for an additional 2 hours (see Methods). Cells were then fixed, subjected IF and then imaged at each indicated time point and conditions. PC-1 (green), TGN46 (red). Scale bar, 10 µm. **E.** Quantification of the experiment in (D) shows the PC-I fluorescence intensity in the cell at each indicated condition. (Data are means ± SEM of at least 3 experiments. **p<0.01, ****p<0.0001 [One-way ANOVA]).

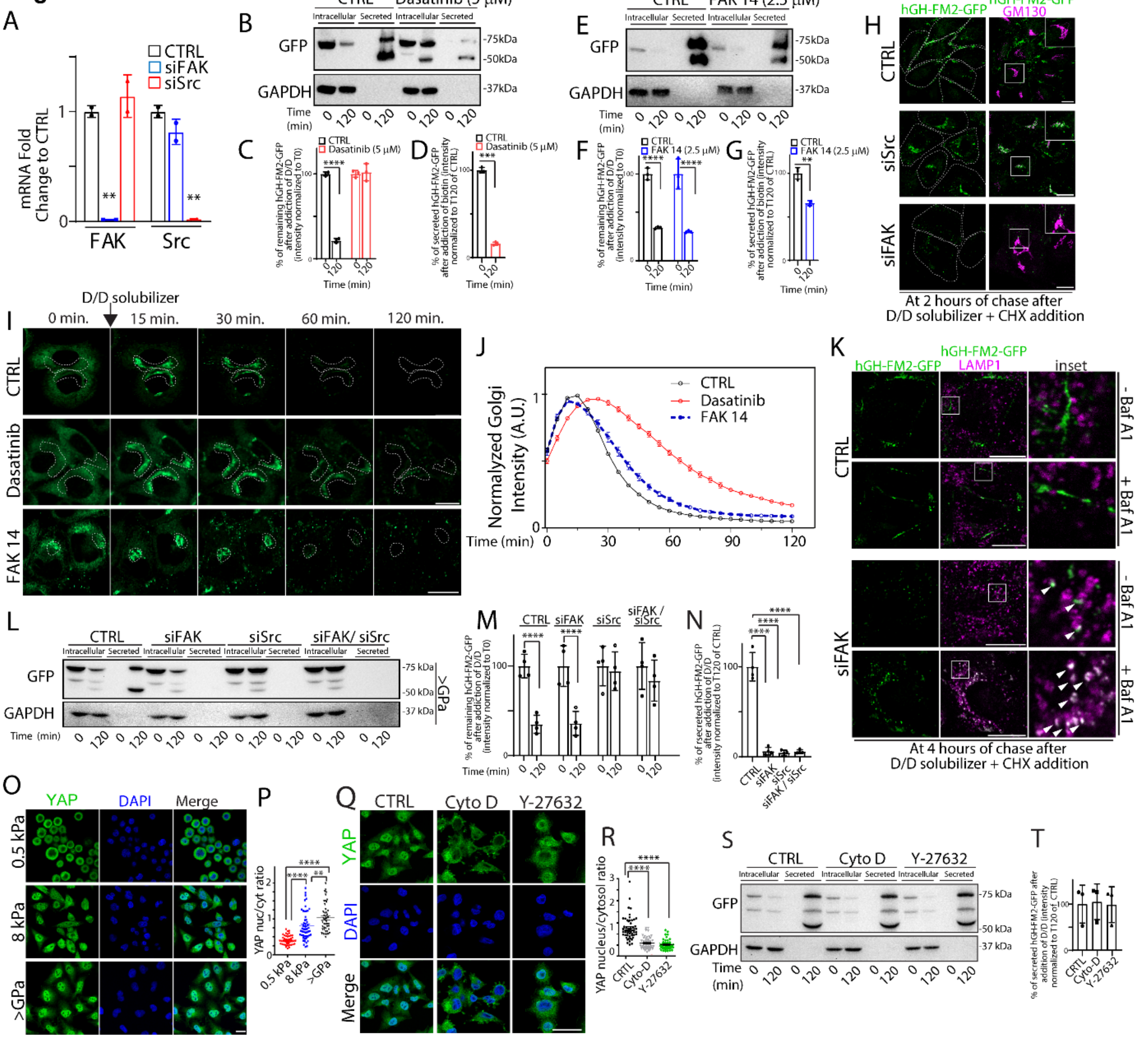


**Figure S4. Effect of matrix stiffness on hGH-FM2-GFP secretion is YAP-independent.**

**A.** mRNA levels of FAK and Src were evaluated in HeLa-GH mock-treated (CTRL) or interfered for FAK (siFAK) and Src (siSrc) respectively by Real-Time PCR (data are means of at least 3 independent experiments ± SD. **p<0.01 [One-way ANOVA]). **B-G**. Upper panels: HeLa-GH cells cultured on >GPa substrate and either mock-treated (CTRL) or treated with Dasatinib (Src inhibitor; 5 µM) (B) or FAK 14 (FAK inhibitor; 2.5 µM) (E) pretreated for 30 minutes and for the whole duration of the experiment. Cells were then treated with D/D (1 µM) and CHX (50 µg/ml) for 2 hours. The cell lysate (intracellular) and the medium (secreted) were collected at indicated times and analysed by SDS-PAGE (20% of the total intracellular lysate and 100% of the total secreted was loaded, respectively) followed by Western blot with the indicated antibodies (n = 3). Bottom panels: densitometric quantification of intracellular (C, F) and secreted (D, G) hGH-FM2-GFP from (B) and (E) after 2 hours is shown (data are mean of at least 3 experiments ± SD. **p<0.01, ***p<0.001 [One-way ANOVA]); ****p<0.0001 [Two-way ANOVA]). **H**. HeLa-GH cells mock-treated (CTRL) or interfered for FAK (siFAK) or Src (siSrc) were grown on >GPa substrate, exposed to D/D (1 μM), CHX (50μg/ml), fixed after 2 hours and processed for IF labelling: hGH-FM2-GFP (green), GM130 (magenta). Dashed lines indicate cells. White boxes indicate the inset. Scale bar, 20 μm. **I.** Analysis of synchronized transport of hGH-FM2-GFP in HeLa cells cultured on >GPa substrate and either mock-treated (CTRL), treated with Dasatinib (Src inhibitor; 5 μM), or FAK 14 (FAK inhibitor; 2.5 μM). Cells were treated with D/D (1 μM) and CHX (50 μg/ml), and the inhibitors were applied for the entire duration of the trafficking period (2 hours). Cells were imaged by live-cell video microscopy. Representative micrographs show cells at the indicated trafficking time points. Dashed lines outline the Golgi region. Scale bar: 20 μm. **J.** Quantification of the experiment in (I) shows the hGH-FM2-GFP fluorescence intensity in the Golgi region at different time points after D/D solubilizer and CHX addition. Intensity normalized to the maximum peak of the Golgi. Data are means ± SEM. **K**. HeLa-GH cells mock-treated (CTRL) or interfered for FAK (siFAK) were grown on >GPa substrate, pre-treated with Baf A1 (20 nM) for 2 hours and then treated with D/D (1 μM) and CHX (50 μg/ml) for 4 hours in the presence of Baf A1 (20 nM), fixed and processed for IF labelling: hGH-FM2-GFP (green), LAMP1 (magenta). Dashed lines indicate cells. White boxes indicate the inset. Arrowheads indicate hGH-FM2-GFP and LAMP1 co-localisation. Scale bar, 20 μm. **L.** HeLa-GH cells mock-treated (CTRL) or silenced for FAK (siFAK), Src (siSrc) or for both the kinases were grown on >GPa substrate. Cells were then treated with D/D (1 μM) and CHX (50 μg/ml) for 2 hours. The cell lysate (intracellular) and the medium (secreted) were collected at indicated times and analysed by SDS-PAGE (50% of the total intracellular lysate and 100% of the total secreted was loaded, respectively) followed by Western blot with the indicated antibodies (n = 3)**.** **M, N.** Densitometric quantification of (L) shows intracellular (M) or secreted (N) hGH-FM2-GFP after 2 hours (data are mean of at least 3 experiments ± SD. ****p<0.001 [One-way ANOVA])**.** **O.** Fluorescence microscopy images of HeLa-GH cells grown on indicated substrate rigidities for 24 hours, fixed and processed for IF labelling: YAP (green), DAPI (blue). Scale bar, 20 µm. **P**. Quantification of the nuclear/cytoplasmic ratio of YAP of the experiment in (O); (individual data points are shown in the graphs; data are means ± SEM of at least 3 independent experiments. **p<0.01, ****p<0.0001 [One-way ANOVA]). **Q.** Representative confocal images of YAP staining of HeLa-GH cells grown on >GPa for 24 hours, and treated with, Cytochalasin D (Cyto D, 1 µM for 16 hours) or Y-27632 (50 µM for 16 hours). Cells were then fixed and processed for IF labelling: YAP (green), DAPI (blue). Scale bar, 150 µm. **R.** Quantification of the nuclear/cytoplasmic ratio of YAP of the experiment in (Q); (individual data points are shown in the graphs; data are means ± SEM of at least 3 independent experiments. ****p<0.0001 [One-way ANOVA]). **S.** Cells were treated as in (Q) and the cell lysate (intracellular) and the medium (secreted) were collected at indicated times and analysed by SDS-PAGE (20% of the total intracellular lysate and 100% of the total secreted was loaded, respectively) followed by Western blot with the indicated antibodies (n = 3). **T**. Densitometric quantification of the secreted hGH-FM2-GFP levels in (S) is plotted (data are means of at least 3 experiments ± SD).

**Figure S5**

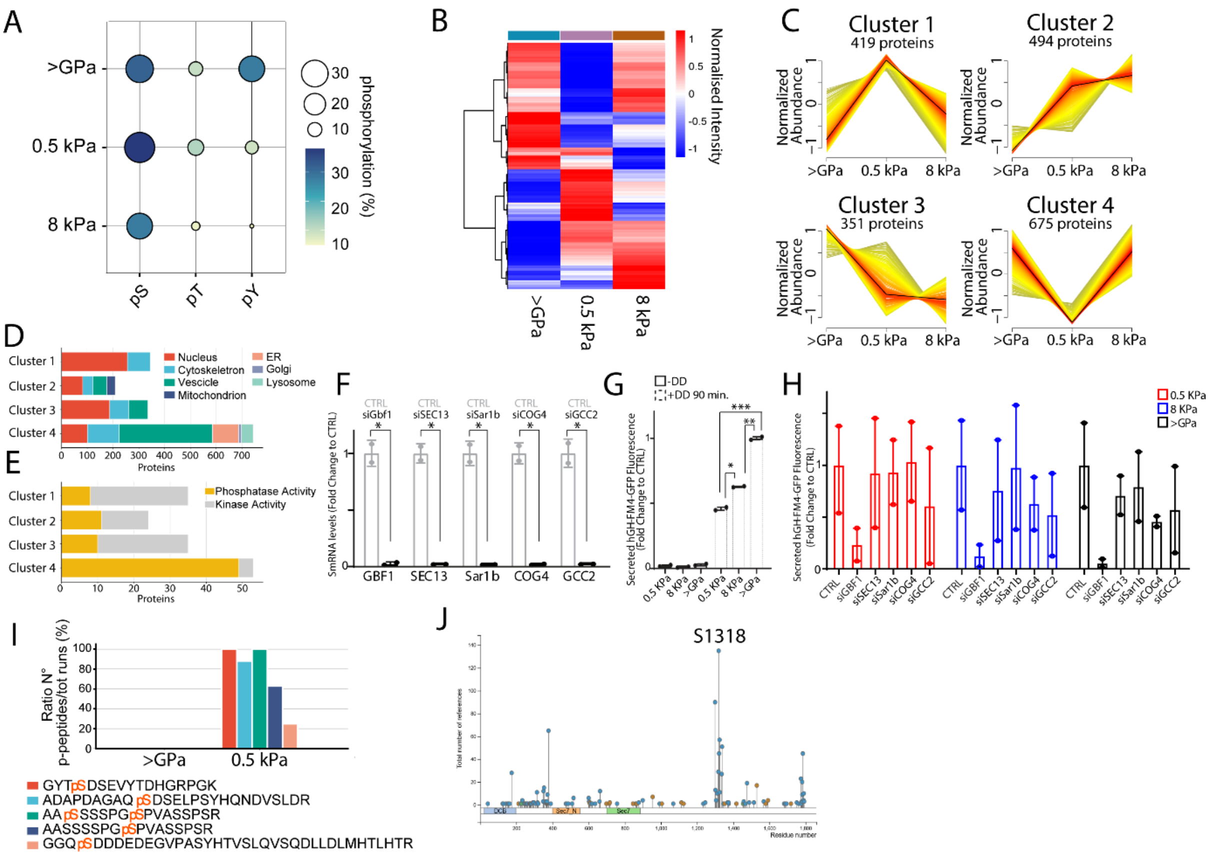

**Figure S5. Influence of stiffness on the phosphorylation proteome of HeLa-GH cells.**

**A.** Chart showing the frequency distribution of phospho-sites, identified in the phosphoproteomics analysis of HeLa-GH cells grown on 0.5 kPa and 8 kPa and >GPa substrates. **B**. Heatmap (Log2 LFQ normalized intensity) of the significantly deregulated phosphoproteins (Anova, FDR <0.05) among the 3 experimental conditions. **C**. Profile plots displaying the LFQ intensities of phosphoproteins grouped into clusters that are significantly different (Anova) among 0.5 kPa and 8 kPa and >GPa groups. **D, E.** Bar plots representing the GOCC and GOMF of the differently regulated phosphoproteins among the 3 experimental conditions. **F**. mRNA levels of Gbf1, SEC13, Sar1b, COG4 and GCC2 were evaluated in HeLa-GH mock-treated (CTRL) or interfered for Gbf1, SEC13, Sar1b, COG4 and GCC2 respectively by Real-Time PCR (data are means ± SD of at least 2 independent experiments. *p<0.05, [Student's t-test]). **G.** HeLa-GH cells were cultured on indicated substrate stiffness.

After 24 hours, cells were treated with D/D (1 μM) and CHX (50 μg/ml) for 2 hours to induce hGH-FM2-GFP secretion. Then, GFP fluorescence of the cell supernatant was measures and normalized to internal DAPI staining (data are means ± SD of at least 2 independent experiments. *p<0.05, **p<0.01, ***p<0.001 [Two-way ANOVA]). **H**. HeLa-GH cells were silenced for the indicated genes for 48 hours and then plated on 2D silicone substrates with indicated substrate stiffness. After 24 hours, cells were treated as in (G) (data are means ± SD of at least 2 independent experiments. **I**. Bar plot depicting the distribution of phospho-peptides associated with GBF1 in >GPa and 0.5 kPa groups. The y-axis shows the number of phospho-peptides normalized to the total MS-run count, while the x-axis represents the different experimental conditions. **J**. Schematic representation of GBF1 phosphorylation revealed by MS analysis.

**Figure S6**

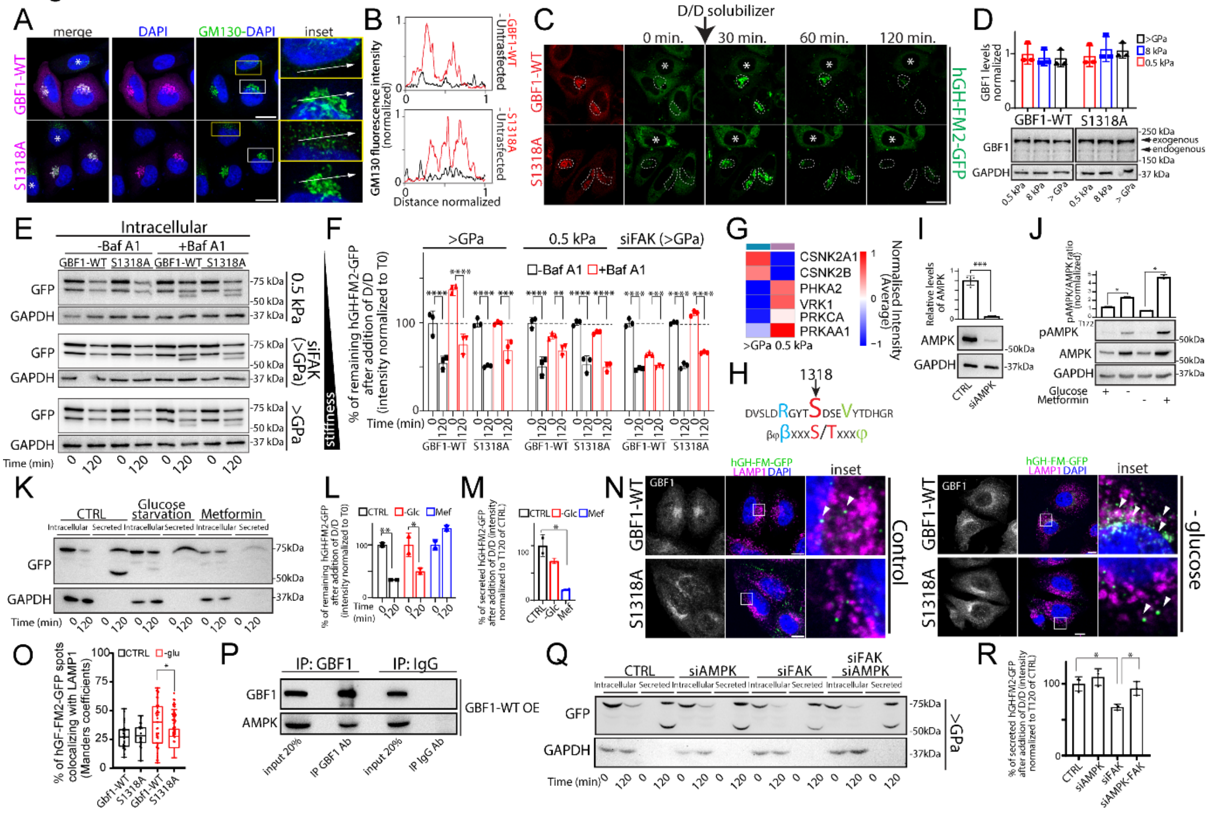

**Figure S6. Phospho-regulation of GBF1 at Ser1318 controls cargo sorting and secretion in response to mechanical cues.**

**A**. Visualization of Golgi morphology in HeLa-GH cells transfected with BFA-resistant GBF1-WT or S1318A mutant, seeded on >GPa substrate, and treated with BFA (0.25 µg/ml) for 2 hours. Cells were fixed and processed for IF labelling: GM130 (green) and GBF1 (magenta). Yellow box indicates the inset of non-transfected cells. White box indicates the inset of transfected cells. Scale bar, 20 µm. **B**. Fluorescence intensity distribution of markers along the line scan (arrow in [A]) for line-scan analyses. **C**. Synchronized transport of hGH-FM2-GFP in HeLa-GH cells expressing BFA-resistant GBF1-WT or S1318A mutant, seeded on >GPa substrate. Cells were then treated with BFA (0.25 µg/ml) for 30 minutes prior to expose them to D/D (1 µM), CHX (50 µg/ml) and BFA (0.25 µg/ml) and then visualised by video microscopy for 2 hours. Micrographs showing cells at the indicated trafficking time points. Dashed lines indicate the Golgi region. Asterisks indicate non-transfected cells. Scale bar, 20 µm. **D**. (Bottom panel) HeLa-GH cells transfected with BFA-resistant GBF1-WT or S1318A mutant, seeded on the indicated substrate for 24 hours, lysed, and subjected to SDS-PAGE and Western blot to analyse transfection levels. Arrows indicate endogenous and exogenous GBF1. (Upper panel) Densitometric quantification of the blot shows the amount of transfected GBF1-WT or S1318A mutant (data are mean of at least 3 experiments ± SD). **E**. Mock-treated (CTRL) or siFAK HeLa-GH cells and expressing BFA-resistant GBF1-WT or S1318A mutant, were seeded on indicated substrates. Cells were treated with BFA (0.25 µg/ml) for 30 minutes prior to expose them to D/D (1 µM), CHX (50 µg/ml) and BFA (0.25 µg/ml) for 2 hours, with or without Baf A1 (20 nM). Cells were lysed and processed for SDS-PAGE and Western blot with the indicated antibodies. Western blot shows intracellular levels of hGH-FM2-GFP at each indicated timepoint. Data are representative of at least 3 independent experiments. **F**. Densitometric quantification of (E) shows intracellular levels of hGH-FM2-GFP (data are mean of at least 3 experiments ± SD. **p<0.01, ***p<0.001, ****p<0.0001 [Two-way ANOVA]). **G**. Heatmap of kinases potentially phosphorylating S1318, based on data from PhosphoSitePlus. Protein intensities are log10-transformed and represented as colours ranging from red to blue. **H.** The sequence surrounding S1318 in the DVSLDRGYTSDSEVYTDHGR peptide represents a consensus motif for AMPK phosphorylation. **I.** HeLa-GH cells mock-treated (CTRL) or silenced for AMPK (siAMPK) were lysed and the cell lysate was analysed by SDS-PAGE followed by Western blot with the indicated antibodies (data are mean of at least 3 experiments ± SD. ***p<0.001, [Student's t-test]). **J.** HeLa-GH cells grown on >GPa substrate (CTRL) were subjected to glucose starvation or Metformin (5 μM) for 16 hours. Cells were then lysed and processed for SDS-PAGE and Western blot with the indicated antibodies (data are mean of at least 3 experiments ± SD. *p<0.05, [Student's t-test]). **K.** HeLa-GH cells treated as in (J) were then exposed to D/D (1 μM), CHX (50 μg/ml) for 2 hours. The cell lysate (intracellular) and the medium (secreted) were collected at indicated times and analysed by SDS-PAGE (20% of the total intracellular lysate and 100% of the total secreted was loaded, respectively) followed by Western blot with the indicated antibodies (n=3). **L, M.** Densitometric quantification of (K) shows intracellular (L) or secreted (M) levels of hGH-FM2-GFP after 2 hours (data are mean of at least 3 experiments ± SD. *p<0.05, **p<0.01 [Two-way ANOVA]). **N.** Synchronized transport of hGH-FM2-GFP in HeLa-GH cells expressing BFA-resistant GBF1-WT or S1318A mutant, seeded on >GPa substrate and subjected to glucose starvation for 16 hours. Cells were then treated with BFA (0.25 μg/ml) for 30 minutes prior to expose them to D/D (1 μM), CHX (50 μg/ml) and BFA (0.25 μg/ml) for 2 hours, fixed and processed for IF labelling: hGH-FM2-GFP (green), GBF1 (white), DAPI (blue) and LAMP1 (magenta). White box indicates the inset. Arrowhead indicates the colocalization of hGH-FM2-GFP and LAMP1. Scale bar, 20 μm. **O.** Quantification of hGH-FM2-GFP colocalization with LAMP1 lysosomal marker from (N). Colocalization was calculated from >20 cells using ImageJ (JACoP plugin; Manders coefficients) (data are mean of at least 3 experiments ± SEM. *$p<0.01$, One-way ANOVA]). **P.** HeLa-GH overexpressing GBF1-WT were grown on 0.5 kPa. Then cell lysates were precipitated using anti-GBF1 and IgG antibodies and the immunoprecipitates were subjected to Western blotting using the indicated antibodies. **Q.** HeLa-GH cells mock-treated (CTRL) or silenced for AMPK (siAMPK), FAK (siFAK) or for both the kinases were grown on >GPa substrate. Cells were then treated with D/D (1 μM) and CHX (50 μg/ml) for 2 hours. The cell lysate (intracellular) and the medium (secreted) were collected at indicated times and analysed by SDS-PAGE (80% of the total intracellular lysate and 100% of the total secreted was loaded, respectively) followed by Western blot with the indicated antibodies (n = 3). **R**. Densitometric quantification of (Q) shows secreted hGH-FM2-GFP after 2 hours (data are mean of at least 3 experiments ± SD. *$p<0.05$ [One-way ANOVA]).

### Figure S7

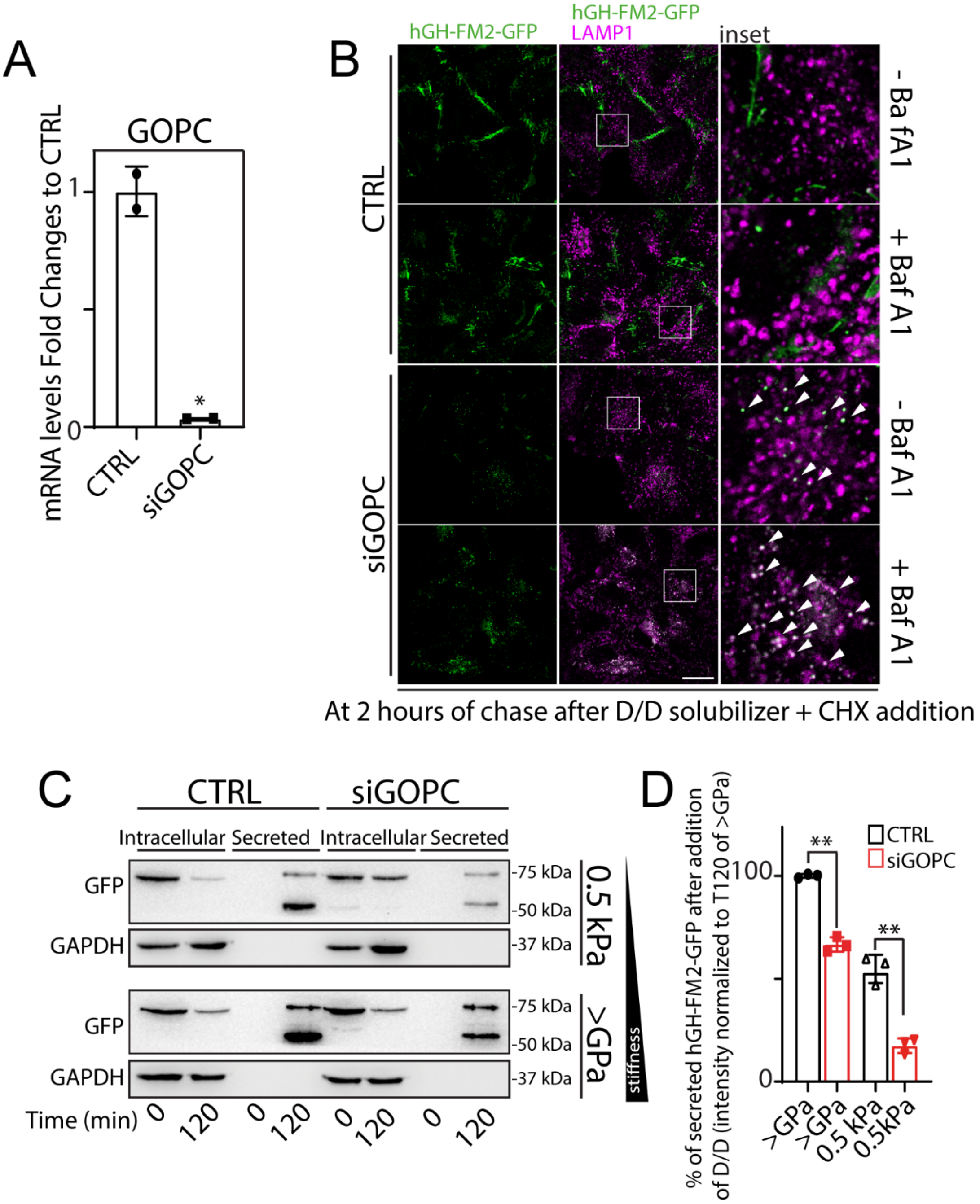


**Figure S7. GOPC silencing affects hGH-FM2-GFP secretion independently of substrate stiffness.**

**A.** GOPC mRNA levels measured by real-time PCR in HeLa-GH cells silenced for GOPC (siGOPC) for 72 hours (data are mean of at least 3 experiments ± SD. *p<0.05, [Student's t-test]). **B**. Visualization of hGH-FM2-GFP transport in HeLa-GH cells mock-treated (CTRL) or silenced for GOPC (siGOPC), grown on >GPa substrate and treated with D/D (1 µM) and

CHX (50 μg/ml) for 2 hours. Fixed cells were processed for IF labelling: hGH-FM2-GFP (green) and LAMP1 (magenta). White boxes indicate the inset. Arrowheads indicate colocalization. Scale bar, 20 μm. **C**. HeLa-GH cells mock-treated or silenced for GOPC (siGOPC) were grown on the indicated substrate. Cells were then treated with D/D (1 μM) and CHX (50 μg/ml) for 2 hours. The cell lysate (intracellular) and the medium (secreted) were collected at indicated times and analysed by SDS-PAGE (20% of the total intracellular lysate and 100% of the total secreted was loaded, respectively) followed by Western blot with the indicated antibodies (n=3). **D**. Densitometric quantification of (C) shows secreted hGH-FM2-GFP after 2 hours (data are mean of at least 3 experiments ± SD. **p<0.01, [Student's t-test]).

**Supplementary Movies legends**

**Movie S1. hGH-FM2-GFP synchronized transport in HeLa-GH grown on indicated substrate rigidities.**

After 24 hours of cell seeding, at time 0, RPMI containing cycloheximide (50 μg/ml) and D/D solubilizer (1 μM) was added to cells. The trafficking of hGH-FM2-GFP was monitored using confocal microscope. Images were acquired at 5 minutes intervals.

**Movie S2. hGH-FM2-GFP synchronized transport in HeLa-GH cells overexpressing GBF1-WT**

HeLa-GH cells were transfected to express GBF1-WT. After 24 hours of expression, cells were treated with BFA (0.25 μg/ml) for 30 minutes. Then, at time 0, CHX (50 μg/ml) and D/D solubilizer (1 μM) was added to cells and the trafficking of hGH-FM2-GFP was monitored using confocal microscope. Images were acquired at 5 minutes intervals. Asterisks indicate non transfected cells (see Figure S6C).

**Movie S3. hGH-FM2-GFP synchronized transport in HeLa-GH cells overexpressing S1318A mutant.**

HeLa-GH cells were transfected to express S1318A mutant. After 24 hours of expression, cells were treated with BFA (0.25 µg/ml) for 30 minutes. Then, at time 0, cycloheximide (50 µg/ml) and D/D solubilizer (1 µM) was added to cells and the trafficking of hGH-FM2-GFP was monitored using confocal microscope. Images were acquired at 5 minutes intervals. Asterisks indicate non transfected cells (see Figure S6C).

**Supplementary tables**

**Table S1.** List of antibodies used in this study.

| **REAGENT** | **SOURCE** | **IDENTIFIER** |
|---|---|---|
| Polyclonal Rabbit GOLPH3 | Abcam | Cat#ab98023 |
| Polyclonal TGN46 antibody | Bio-Rad | Cat#AHP500GT |
| Monoclonal Rabbit AMPK alpha 1 | Abcam | Cat#ab32047 |
| Monoclonal Rabbit Phospho-AMPK T172 | Abcam | Cat#ab314032 |

| | | |
|---|---|---|
| Monoclonal Mouse GAPDH | Santa Cruz Biotechnology | Clone 6C5<br>Cat #sc-32233 |
| GFP | Abcam | Cat#ab290 |
| Polyclonal Rabbit LAMP1 | Abcam | Cat#ab24170 |
| Polyclonal COL1A1 | Invitrogen | Cat #PA5-29569 |
| YAP (D8H1X) XP | Cell Signalling Technologies | Cat #14074 |
| GBF1 | Abcam | Cat #86071 |
| Monoclonal Mouse GM130 | BD Biosciences | Clone 35<br>Cat # 610822 |
| Monoclonal Mouse anti-β Actin | Sigma | Clone AC-74<br>Cat #A2228 |
| Polyclonal Rabbit B4GALT1 | Sigma | Cat # HPA010807 |
| Monoclonal Mouse GRASP65 | Santa Cruz Biotechnology | Clone D12<br>Cat # sc-374423 |
| Monoclonal Mouse Sec31A | BD Biosciences | Clone 32 Cat #612351 |

| | | |
|---|---|---|
| Monoclonal Mouse FAK | Santa Cruz Biotechnology | CloneH1<br>Cat # sc-1688 |
| Polyclonal phospho-FAK Y397 | Invitrogen | Cat #PA5-17084 |
| Polyclonal Rabbit SRC | Invitrogen | Cat # 44-655G |
| Polyclonal phospho-SRC Y418 | Invitrogen | Cat # 44-660G |
| Anti-rabbit, donkey Alexa Fluor 488 | ThermoFisher Scientific | Cat#A-21206 |
| Anti-rabbit, donkey Alexa Fluor 568 | ThermoFisher Scientific | Cat#A10042 |
| Anti-mouse, donkey Alexa Fluor 568 | ThermoFisher Scientific | Cat#A10037 |
| Anti-mouse, donkey Alexa Fluor 488 | ThermoFisher Scientific | Cat#A-21202 |
| Anti-rabbit, goat IgG H & L peroxidase conjugated | Millipore | Cat# 401315 |
| Anti-mouse, goat IgG H & L peroxidase conjugated | Millipore | Cat# 401215 |

**Table S2.** List of siRNAs used in this study

| Human Gene | Accession Number | siRNA Sequence |
|---|---|---|
| Src | NM_005417 | #1: 5'- GAGAACCUGGUGUGCAAAG-3'<br>#2: 5'- CGUCCAAGCCGCAGACUCA-3'<br>#3: 5'- CCUCAGGCAUGGCGUACGU-3' |
| FAK | NM_005607 | #1: 5′-GGUUCAAGCUGGAUUAUUU-3'<br>#2: 5′-CCGGTCGAATGATAAGGTGTA-3′<br>#3: 5'- GGAAAUACAGUUUGGAUCU-3' |
| GOPC | NM_020399 | #1: 5'-GGCCUUGGCAUUUCAAUUA-3'<br>#2: 5'-GGACAUCGUUACCGUUUGU-3' |

| | | |
|---|---|---|
| GCC2 | NM_181453 | #1: 5'-UCAAAGAGAUACCAUGUUA-3'<br>#2: 5'-GAGCAGAGUUGAUACUAUU-3' |
| Sar1B | NM_016103 | #1: 5'-CGACAGACCUGAAGCCAUC-3'<br>#2: 5'-CGAGAGAUGUUUGGUUUAU-3' |
| COG4 | NM_015386 | #1: 5'-UCUAUACCCUGAUCAAAUA-3'<br>#2: 5'-GCUGGAGGCUGUAUACGAA-3' |
| GBF1 | NM_004193 | #1: 5'-GUCCAUCCCUGAAGUGUUA-3'<br>#2: 5'-GCAAGGACUUUGAGCAAGA-3' |
| SEC13 | NM_001136026.3 | #1: 5'-GCUCAAGACUGCUAGAAAU-3'<br>#2: 5'-CCACAAAGCUCAGUCAAAA-3' |
| AMPK | NM_001139676 | #1: 5'-CAAAGUCGACCAAAUGAUA-3' |

| | | |
|---|---|---|
| | | #2: 5’-GUAGAGCAAUCAAACAAUU-3’ |
| AllStars Negative Control siRNA | Qiagen | Cat #SI03650318 |

**Table S3.** Primers used in this study to determine mRNA levels of indicated gene

| **Human** | **Forward Primer** | **Reverse Primer** |
|---|---|---|
| IL6 | 5’-CCACTCACCTCTTCAGAACG-3’ | 5’-CATCTTTGGAAGGTTCAGGTTG-3’ |
| ANG | 5’-TGTTGGAAGAGATGGTGATGG-3’ | 5’-CATAGTGCTGGGTCAGGAAG-3’ |
| HPRT1 | 5’-AGCTTGCTGGTGAAAAGGAC-3’ | 5’-GTCAAGGGCATATCCAACAAC-3’ |
| FGF6 | 5’-GGTGAGTCTCTTTGGAGTGAG-3’ | 5’-AAGTCTGACTCGTAGGCATTG-3’ |

| | | |
|---|---|---|
| TIMP1 | 5'-TTCTGCAATTCCGACCTCG-3' | 5'-TCATAACGCTGGTATAAGGTGG-3' |
| TIMP2 | 5'-CCCTCTGTGACTTCATCGTG-3' | 5'-GAGATGTAGCACGGGATCATG-3' |
| IGFBP3 | 5'-CAGAGCACAGATACCCAGAAC-3' | 5'-AGCACATTGAGGAACTTCAGG-3' |
| IL12B | 5'-CACATTCCTACTTCTCCCTGAC-3' | 5'-CTGAGGTCTTGTCCGTGAAG-3' |
| PLGF | 5'-ACATGTTCAGCCCATCCTG-3' | 5'-GTCCCCAGAACGGATCTTTAG-3' |
| TGFB2 | 5'-CCCCACATCTCCTGCTAATG-3' | 5'-ATGTAAAGTGGACGTAGGCAG-3' |
| TGFB3 | 5'-AGTGGCTGTCCTTTGATGTC-3' | 5'-CACCTCGTGAATGTTTTCCAG-3' |
| IGFBP2 | 5'-ACATCCCCAACTGTGACAAG-3' | 5'-ATCAGCTTCCCGGTGTTG-3' |
| CXCL1 | 5'-AACCGAAGTCATAGCCACAC-3' | 5'-CCTCCCTTCTGGTCAGTTG-3' |

| | | |
|---|---|---|
| CXCL5 | 5’-TCTGCAAGTGTTCGCCATAG-3’ | 5’-CAGTTTTCCTTGTTTCCACCG-3’ |
| NT3 | 5’-GATGCCATGGTTACTTTTGCC-3’ | 5’-AATGAGGGAATTGAGCGAGTC-3’ |
| G-CSF | 5’-TTCCTGCTCAAGTGCTTAGAG-3’ | 5’-AGCTTGTAGGTGGCACAC-3’ |
| Src | 5’-CAATGCAGAGAACCCGAGAG-3’ | 5’-TGCGGATCTTGTAGTGCTTC-3’ |
| FAK | 5’-AAATACGGCGATCATACTGGG-3’ | 5’-TTGGCCTTGACAGAATCCAG-3’ |
| GOPC | 5’-GTTGAGAAAAGAGAATGAAGCCC-3’ | 5’-TCCCTTCATATCTCGTCCTAGC-3’ |
| GCC2 | 5’-TTCAGAAGCTCAGTTCCACC-3’ | 5’-TCTCGCTCTTGATTCCTTTCC-3’ |
| SEC13 | 5’-TGCAGCAGATTGAGGAGGAC-3’ | 5’-CCTCCTGCTGTTGTTGTTGG-3’ |

| | | |
|---|---|---|
| Sar1B | 5'-TCCCAACATTACATCCCACTTC-3' | 5'-GCCATTGATAGCAGGAAGGTAG-3' |
| COG4 | 5'-TTTTCTCCGTCTCCCACAAC-3' | 5'-CTTGAACTCTGCCATTTGCTG-3' |
| GBF1 | 5'-CTGCTTCCCTGCGAGTATG-3' | 5'-CCAGTGCCATCTCCTTCATC-3' |

**Table S4.** All the commercial kits used for cDNA extraction and mRNA analysis.

| Commercial assays and kits | | |
|---|---|---|
| RNA easy mini | Qiagen | Cat #74106 |
| QuantiTect Reverse Transcription Kit | Qiagen | Cat #205311 |
| SYBR™ Green PCR Master Mix | ThermoFisher Scientific | Cat #4309155 |
